# Maternal immune activation and peripubertal stress differentially disrupt glutamatergic, endocannabinoid, and neuromodulatory signalling in the adult rat dorsal hippocampus: implications for excitatory-inhibitory balance

**DOI:** 10.64898/2026.06.13.732070

**Authors:** Pablo C. del Olmo, Charlene Nowotny, Mario Moreno-Fernández, Roberto Capellán, Javier Orihuel, Alberto Marcos, Emilio Ambrosio, Marcos Ucha, Alejandro Higuera-Matas

**Author notes:** **Corresponding author:** Alejandro Higuera-Matas, PhD, Department of Psychobiology. School of Psychology. National University for Distance Learning (UNED). C/Juan del Rosal 10. Madrid, 28040. Spain. Melanoma Laboratory, Molecular Oncology Programme, Spanish National Cancer Research Centre (CNIO), Madrid, Spain. Laboratory of Neuropharmacology-Neurophar, Department of Medicine and Life Sciences, Universitat Pompeu Fabra (UPF), Barcelona, Spain. Neurobiology of behavior Research Group, Department of Medicine and Life Sciences, Universitat Pompeu Fabra (UPF), Barcelona, Spain.

## Abstract

Disruptions in excitatory–inhibitory (E/I) balance during neurodevelopment have been implicated in a range of psychiatric conditions, yet the neurochemical alterations associated to early-life insults and their potential contribution to E/I imbalance remain poorly understood. Using a “two-hit” rat model combining maternal immune activation (MIA; lipopolysaccharide -LPS- on gestational days 15–16) and peripubertal unpredictable stress (PUS; postnatal days 28–38), we examined the long-term effects of these insults, alone and in combination, on the adult dorsal hippocampus. Assessments included gene and/or protein expression of glutamatergic and GABAergic markers, endocannabinoid system enzymes, neuromodulatory amino acid level and prepulse inhibition (PPI) of the acoustic startle response. MIA increased GluN1 protein expression, while PUS reduced the *Grin2a*/*Grin2b* mRNA ratio, indicating incomplete NMDA receptor subunit maturation. GABA levels and GABA-Aγ2 expression were unchanged, suggesting deficient inhibitory compensation in the face of heightened excitatory tone. PUS increased *Mgll* gene expression, whereas a trend towards reduced *Dagla* expression was observed exclusively in non-stressed LPS-exposed animals, suggesting that MIA may suppress 2-AG synthesis only in the absence of subsequent stress. MIA and PUS displayed interactive effects on taurine levels, with elevation observed only in the double-hit condition; glycine was elevated by MIA independently of PUS. These findings support a model in which MIA and PUS converge on hippocampal E/I balance through complementary adaptations — excitatory upregulation, incomplete synaptic maturation, and reduced endocannabinoid tone — inadequately counterbalanced by inhibitory systems. Taurine and glycine emerge as potential markers of homeostatic compensation in response to early neurochemical dysregulation.

## 1. Introduction

The balance between excitatory and inhibitory (E/I) neurotransmission is essential for normal brain function. In the mammalian brain, excitatory glutamatergic drive and inhibitory GABAergic control must be precisely coordinated to sustain information processing, synaptic plasticity, and cognitive performance (Sears and Hewett, 2021; Sohal and Rubenstein, 2019). This balance is actively regulated throughout neurodevelopment: a key maturational landmark is the progressive switch in NMDA receptor composition from GluN2B- to GluN2A-containing subunits, which reflects the stabilisation of excitatory synapses during late postnatal and adolescent development (Paoletti et al., 2013; Yashiro and Philpot, 2008). Disruption of this trajectory results in sustained excitatory hyperfunction and insufficient inhibitory compensation — a state increasingly recognised as a core feature of schizophrenia and related neurodevelopmental disorders (Rubenstein and Merzenich, 2003; Sohal and Rubenstein, 2019; Uliana et al., 2024). In schizophrenia, hippocampal hyperexcitability driven by reduced parvalbumin-expressing GABAergic interneurons (Lewis et al., 2012; Konradi et al., 2011) and altered NMDA receptor signalling is a particularly robust finding, with downstream consequences for dopaminergic regulation and cognition (Lisman et al., 2008; Konradi et al., 2011; Boley, Perez and Lodge, 2014).

The hippocampus is a central node in the neural circuits disrupted in schizophrenia. In rodents, the dorsal hippocampus — the subregion examined in the present study — is preferentially involved in spatial learning, episodic memory, and contextual fear conditioning, while the ventral portion is more closely associated with emotional and stress-related processing (Moser and Moser, 1998; Fanselow and Dong, 2010). Of note, the previously mentioned cognitive functions are among those most consistently impaired in schizophrenia patients and in animal models of the disorder (Heckers and Konradi, 2015).

Maternal immune activation (MIA), typically modelled by gestational administration of the bacterial endotoxin lipopolysaccharide (LPS), and peripubertal unpredictable stress (PUS) are two environmental insults that may act synergistically to precipitate schizophrenia-relevant phenotypes —a possibility which has come to be known as the “two-hit” hypothesis (Giovanoli et al., 2013; Meyer, 2019).

MIA on one hand has been shown to disrupt hippocampal synaptic properties and information processing in adult offspring (Ito et al., 2010). Indeed, prenatal LPS exposure impairs spatial memory (Batinić et al., 2016) altering NMDA receptor function (Burt et al., 2013; Escobar et al., 2011). More broadly, prenatal immune activation with viral mimetics (such as Poly I:C) has been shown to alter hippocampal excitatory and inhibitory synaptic transmission in offspring, with decreased excitatory and increased GABAergic transmission in CA1 pyramidal cells across development (Nakagawa et al., 2020).

PUS on the other hand, produces spatial memory deficits linked to aberrant hippocampal neurogenesis, synaptic plasticity, and dysregulation of the HPA axis (McCormick et al., 2012; Borsini et al., 2023). In addition, early stress exposure alters the developmental maturation of NMDA receptor subunit composition (Rodenas-Ruano et al., 2012) and disrupts hippocampal endocannabinoid signalling (Zhang et al., 2015).

With regard to the potential synergistic actions of both hits, MIA has been shown to induce enduring alterations in hippocampal glutamatergic signalling, including NMDA receptor hypofunction under basal conditions that is paradoxically sensitised by subsequent stress (Burt et al., 2013), as well as region- and timing-dependent reductions in GABAergic interneuron markers (Gillespie et al., 2024; Giovanoli et al., 2014).

Additional neuromodulatory systems contribute to hippocampal E/I regulation: glycine and taurine act as NMDA receptor co-agonist and inhibitory neuromodulator, respectively (Balu, 2016; Wu and Prentice, 2010), while neuronal nitric oxide synthase (nNOS), physically coupled to the NMDA receptor via PSD-95 scaffolding, modulates both glutamatergic and GABAergic release as a retrograde messenger (Hardingham et al., 2013; Tricoire and Vitalis, 2012). NOS1 — the gene encoding nNOS — is a schizophrenia risk gene (Nasyrova et al., 2015; Freudenberg et al., 2015), and MIA transiently increases nNOS immunoreactivity in neonatal hippocampal subregions (Zhang et al., 2017); however, whether these effects persist into adulthood in the dorsal hippocampus remains unclear.

Previous work from our group has shown that MIA and PUS interact to produce lasting metabolic and glutamatergic alterations in corticostriatal circuits (Capellán et al., 2022, 2023), but their combined impact on hippocampal E/I neurochemistry has not been directly examined. In the present study, we combined RT-qPCR, western blotting, and capillary electrophoresis to characterise the long-term effects of maternal LPS-induced MIA and PUS, alone and in combination, on markers of glutamatergic and GABAergic signalling, the endocannabinoid system, nNOS protein expression, and neuromodulatory amino acids in the adult rat dorsal hippocampus. We hypothesised that MIA and PUS would independently and interactively disrupt hippocampal E/I balance through complementary pathways, and that their combination would unmask neurochemical alterations not evident with either insult alone. Consistent with this hypothesis, we found that MIA increased GluN1 AMPA glutamate receptor subunit protein expression and elevated hippocampal glycine levels, while PUS reduced the GluN2A/GluN2B ratio — indicative of incomplete NMDA receptor maturation — and upregulated MAGL (monoacylglycerol lipase, the main 2-AG degrading enzyme) gene expression. MIA and PUS displayed interactive effects on taurine levels, with the double-hit condition showing an elevation that was not observed with either insult alone. These findings support a model in which MIA and PUS converge on the hippocampal E/I system through distinct but complementary pathways, with neuromodulatory amino acids emerging as potential markers of homeostatic compensation.

## 2. Materials and Methods

### 2.1 Animals

Experiments were performed on the male offspring of 14-week-old male and 12-week-old female Sprague-Dawley rats obtained from Charles River (France). Animals were housed under controlled conditions (temperature 21°C, relative humidity 50–60%, simulated 12 h/12 h light/dark cycle) with ad libitum access to standard rodent chow (A04, Panlab, Barcelona, Spain) and water. Animals were accommodated in polycarbonate cages (48.3 × 26.7 × 20.3 cm). All experimental procedures complied with the guidelines of the European Union for the care of laboratory animals (EU Directive 2010/63/EU) and were authorised by the Autonomous Community of Madrid under procedure number PROEX 078/18 and approved by the UNED Bioethics Committee.

A total of 35 male offspring were used: saline (SAL)/no stress (NS) n = 10, SAL/stress (S) n = 10, LPS/NS n = 8, LPS/S n = 7. Animals were derived from 8 saline-treated litters and 7 LPS-treated litters, with no more than two to three animals per litter assigned to the same experimental group, to prevent litter effects from confounding the statistical analysis.

### 2.2 Experimental Procedure

After a 7-day acclimation period, male and female rats were paired for mating. Pregnancy was confirmed by daily vaginal smears; the presence of sperm was designated as gestational day (GD) 0. On GD 15 and GD 16, pregnant dams received intraperitoneal injections of lipopolysaccharide (LPS; serotype O111:B4, Sigma-Aldrich) dissolved in sterile saline and administered at a dose of 100 µg/kg in a volume of 1 ml/kg body weight. This dose was selected on the basis of prior work demonstrating that it does not significantly compromise dam survival and produces only minimal effects on prepulse inhibition, thereby preserving sensitivity to potential unmasking effects of later stressors. Previous studies from our group using the same protocol confirmed that LPS administration at this dose induces transient hypothermia and body weight alterations in the dams (Santos-Toscano et al., 2016; Capellán et al., 2022). Control dams received an equivalent volume of sterile saline on the same days. Dams were randomly assigned to LPS or control conditions. To standardise litter size and reduce variability due to maternal behaviour, each litter was culled to 12 pups at birth. At postnatal day (PND) 21, offspring were weaned and housed in groups of three to four per cage. Between PND 28 and 38, male offspring assigned to the peripubertal stress (PUS) condition were exposed to an unpredictable stress protocol adapted from Giovanoli et al. (2013), consisting of five different stressors applied on alternating days in a counterbalanced order: (1) forced swim (10 min in a tank, 40 cm height × 18 cm diameter, water depth 30 cm, 22 ± 1°C); (2) home cage disruption (five bedding changes during the dark phase); (3) agitation stress (30 min on an orbital shaker at 100 rpm); (4) immobilisation (45 min in a cylindrical restrainer under bright light); and (5) water deprivation for 16 h. Control animals were handled by the same experimenter on the same days to control for handling-related arousal (see Capellán et al., 2022, 2023 for further details).

### 2.3 Prepulse Inhibition of the acoustic startle response (PPI)

On PND 70–73, male offspring were subjected to prepulse inhibition (PPI) testing for the first time, conducted in a non-restrictive polycarbonate chamber (28 × 15 × 17 cm) equipped with a vibration-sensitive platform (Cibertec, Madrid, Spain). Final sample sizes for each condition were as follows — PPI4_30: SAL/NS n = 4, SAL/S n = 7, LPS/NS n = 5, LPS/S n = 5; PPI4_120: SAL/NS n = 7, SAL/S n = 8, LPS/NS n = 6, LPS/S n = 6; PPI12_30: SAL/NS n = 9, SAL/S n = 10, LPS/NS n = 8, LPS/S n = 7; PPI12_120: SAL/NS n = 9, SAL/S n = 9, LPS/NS n = 6, LPS/S n = 7. Variation in sample sizes across conditions reflects the exclusion of individual trials with negative PPI values.

Animals were first acclimated for 7 min to a continuous 65 dB background noise, followed by six pulse-alone trials to habituate the startle response. Each session comprised 35 trials presented with a mean inter-trial interval of approximately 15 s: ten pulse-alone trials (120 dB, 40 ms), five null trials (background noise only), and twenty prepulse-plus-pulse trials in which the pulse was preceded by a prepulse of either 69 dB (4 dB above background) or 77 dB (12 dB above background), with stimulus onset asynchronies of 30 ms or 120 ms, yielding four conditions (PPI4_30, PPI4_120, PPI12_30, PPI12_120). Each session lasted approximately 20 min. PPI was calculated as:

*%PPI = [1 − (mean startle amplitude on prepulse+pulse trials / mean startle amplitude on pulse-alone trials)] × 100*

where both means were computed across all trials of each type within the session. Negative values, indicative of startle facilitation, were excluded from the analysis. Negative PPI values, indicative of startle facilitation rather than inhibition, were excluded from the main analysis. To verify that their distribution did not differ systematically across experimental groups — which would suggest a treatment-related effect on prepulse facilitation — the proportion of negative values per group and condition was examined using Χ^2^ tests of independence. No significant differences in facilitation rates were found between MIA and PUS groups in any condition (all p > 0.24), confirming that their exclusion did not introduce a systematic bias.

### 2.4 Hippocampal extraction and homogenization

On postnatal day (PND) 90, animals were sacrificed by quick decapitation under light isoflurane anaesthesia. The dorsal hippocampus was rapidly dissected on ice using standard anatomical landmarks, and both hemispheres were collected. Hemisphere assignment to molecular analysis was counterbalanced across animals: one hemisphere was dedicated to RNA extraction and gene expression analyses, and the contralateral hemisphere to protein and aminoacid quantification, with the allocation of left or right hemisphere to each analytical pipeline randomised across subjects. Samples were snap-frozen and stored at −80°C until analysis. For protein and amino acid quantification, tissue was homogenised in ice-cold buffer containing 50 mM HEPES (pH 7.5), 320 mM sucrose, protease inhibitor cocktail (cOmplete™, Roche), phosphatase inhibitors (PhosSTOP™, Roche), and 20 mM sodium butyrate, using a pellet pestle (Sigma-Aldrich, Z359971).

### 2.5 RT-qPCR

RNA was extracted from dorsal hippocampal tissue using QIAzol™ lysis reagent (Qiagen), and 469 ng of total RNA were reverse-transcribed into cDNA using the iScript™ cDNA Synthesis Kit (Bio-Rad). Quantitative real-time PCR (RT-qPCR) was performed on a CFX96 Touch™ Real-Time PCR Detection System (Bio-Rad) using SsoAdvanced™ Universal SYBR® Green Supermix (Bio-Rad). Each sample was run in duplicate; no-RT and no-template controls were included as negative controls in each plate. Duplicates with a Cq discrepancy exceeding 0.5 cycles or showing abnormal amplification curves were excluded. Product specificity was confirmed by melting curve analysis. The following genes were quantified: glutamatergic system — *Gria1* (GluA1), *Gria2* (GluA2), *Grm1* (mGluR1), *Grin1* (GluN1), *Grin2a* (GluN2A), *Grin2b* (GluN2B); GABAergic system — *Gabrg2* (GABA-Aγ2); endocannabinoid system — *Cnr1* (CB1), *Dagla* (DAGLα), *Mgll* (MAGL), *Napepld* (NAPE-PLD), *Faah* (FAAH). Primer sequences are provided in Table S1. *Gapdh* was used as the reference gene; its expression stability across experimental conditions was verified prior to normalisation. Relative gene expression was calculated using the ΔΔCt method (Pfaffl, 2001), and reaction efficiencies were determined with LinRegPCR software (Ruijter et al., 2009).

### 2.6 Capillary Electrophoresis

Neuromodulatory amino acid concentrations (GABA, L-glutamate, L-glutamine, glycine, taurine, L-serine, and L-aspartate) were quantified in dorsal hippocampal homogenates by Capillary Electrophoresis with Laser-Induced Fluorescence detection (CE-LIF), using a validated method described in detail by Lorenzo et al. (2013). Analyses were performed on a PA 800 Plus apparatus (Beckman Coulter Inc.) equipped with a fused silica capillary (60.2 cm total length, 50 cm effective length and 75 µm internal diameter). Derivatisation was carried out by incubating samples with the fluorescent reagent NBD-F (4-fluoro-7-nitrobenzofurazan) in alkaline borate buffer (10 mM, pH 9.0) with methanol at 60°C for 17 min. Electrophoretic separation was performed under a high-voltage electric field (+21 kV) using 90 mM borate running buffer (pH 10.25) containing 12.5 mM β-cyclodextrin. Detection was carried out by LIF at an excitation wavelength of 488 nm. Amino acids were identified by their characteristic migration times, and peak areas were calculated using 32 Karat™ software (Beckman Coulter). Aminoadipic acid was added to all samples and standards as an internal standard to correct for inter-run variability. Calibration curves were obtained by least-squares regression using eight-point standard series for each analyte, plotting the corrected!zlarea!zlamino!zlacids/corrected!zlarea!zlIS versus concentration.

### 2.7 Protein Quantification

Protein levels of GluN1 and nNOS were assessed by Western blotting. Prior to electrophoresis, total protein concentration in each homogenate was determined using the DC™ Protein Assay (Bio-Rad), and 6.2 µg of total protein per sample were loaded onto 4–15% Mini-PROTEAN® TGX™ Precast Protein Gels (Bio-Rad) for SDS-PAGE separation under reducing conditions. Proteins were then transferred onto low-fluorescence polyvinylidene difluoride (PVDF) membranes (Bio-Rad) using a Trans-Blot® Turbo™ Transfer System (Bio-Rad). To allow normalisation by total protein load, membranes were imaged before blocking using a Amersham Imager 600 (GE Healthcare) with Stain-Free™ technology. Membranes were subsequently blocked in 1% BSA in Tris-buffered saline with 0.1% Tween-20 (TBS-T) for 1 h at room temperature, and incubated overnight at 4°C with primary antibodies diluted in blocking solution. GluN1 was detected using a rabbit anti-GluN1 antibody (Cell Signaling Technology, cat. no. D65B7; expected molecular weight ∼120 kDa; observed band: 120 kDa; see Figure S1) and nNOS using a rabbit anti-nNOS antibody (Cell Signaling Technology, cat. no. C7D7; expected molecular weight ∼160 kDa; observed band: 150 kDa) (See Figure S1). Sequential detection was performed on the same membrane: after imaging the GluN1 signal, membranes were stripped using a stripping buffer (Abcam: https://www.abcam.com/en-us/technical-resources/guides/western-blot-guide/membrane-stripping-for-western-blot) and re-probed with the anti-nNOS antibody following the same blocking and incubation procedure. After primary antibody incubation, membranes were washed in TBS-T and incubated for 1 h at room temperature with a horseradish peroxidase-conjugated anti-rabbit secondary antibody (Abcam; cat. no. ab6721). Immunoreactive bands were visualised by enhanced chemiluminescence (ECL) and imaged on the Amersham Imager 600. Band intensities were quantified using ImageJ software (National Institutes of Health, Bethesda, MD, USA) and normalised to the corresponding total protein signal from the Stain-Free™ image of the same membrane.

### 2.8 Statistical Analysis

Data analysis was conducted using IBM SPSS Statistics 27 for Windows, with the significance level set at α = 0.05. Prior to analysis, normality of the distribution of each variable was assessed using the Shapiro-Wilk test. Outliers were identified and removed using the interquartile range criterion (IQR × 3). The effects of MIA (prenatal treatment: saline vs. LPS) and PUS (postnatal treatment: no stress vs. stress) on each dependent variable were examined using two-way analysis of variance (ANOVA), with MIA and PUS as between-subjects factors. When a statistically significant interaction between MIA and PUS was detected, simple effects analyses were conducted as follow-up tests, with p-values adjusted using the Bonferroni correction for multiple comparisons. Non-significant trends are reported when F-values approached but did not reach the significance threshold (0.05 < p < 0.10). Data are expressed as mean ± standard error of the mean (SEM). Detailed descriptive and inferential statistics for all variables are provided in the Supplementary Information.

PPI data were analysed using linear mixed models (LMM) implemented in Python (statsmodels 0.14), to exploit the within-subject structure of the four PPI conditions. In this framework, MIA and PUS were entered as between-subjects fixed factors, and prepulse intensity (dB above background: 4 or 12 dB) and stimulus onset asynchrony (ISI: 30 or 120 ms) were entered as within-subject fixed factors, along with all two-way, three-way, and four-way interactions. A random intercept per subject was included to account for individual differences in baseline PPI and the non-independence of repeated observations within the same animal. Missing values arising from the exclusion of negative PPI scores were handled by restricted maximum likelihood estimation (REML), which uses all available observations per subject without requiring listwise deletion. Separate models were fitted for each level of dB (4 dB and 12 dB subsets) and each level of ISI (30 ms and 120 ms subsets) to facilitate interpretation of interactions. The proportion of negative PPI values (prepulse facilitation) was compared across groups using chi-square tests of independence. The following assumptions of the LMM were verified for the 12 dB model as representative: normality of residuals (Shapiro-Wilk: W = 0.984, p = 0.563), homogeneity of variance across groups (Levene: F(3,60) = 0.270, p = 0.847), absence of influential outliers (no standardised residual exceeded ±2.5), and normality of random effects (Shapiro-Wilk on subject intercepts: W = 0.965, p = 0.346). All assumptions were satisfied. Full LMM results are provided in the Supplementary Information.

## 3. Results

### 3.1 Prepulse inhibition

Descriptive statistics for all PPI conditions are provided in Supplementary Table S2. Their distribution did not differ significantly between MIA and PUS groups in any condition (chi-square tests, all p > 0.24), confirming that their exclusion did not introduce a systematic group bias.

The PPI data are presented in Figure 1. Standard factorial ANOVA data are provided in Table S3. With regard to LLM results, in the model restricted to 12 dB prepulse trials, PUS produced a significant increase in PPI regardless of ISI (main effect of PUS: β = 15.55, SE = 7.04, p = 0.027), indicating a consistent facilitating effect of peripubertal stress on sensorimotor gating at high prepulse intensities. A non-significant trend towards a MIA × PUS interaction was also observed at 12 dB (p = 0.085), suggesting that LPS exposure may attenuate the PUS-induced facilitation of PPI. In the model restricted to 4 dB prepulse trials, a significant PUS × ISI interaction was detected (p = 0.044): PUS increased PPI at ISI 30 ms but reduced it at ISI 120 ms, indicating that the direction of the PUS effect at low prepulse intensities is dependent on the temporal gap between prepulse and pulse. When models were fitted separately by ISI, a significant PUS × dB interaction emerged at ISI 120 ms (p = 0.001), reflecting the opposing effects of PUS at 4 dB and 12 dB. No significant effects of MIA were detected in any LMM (all p > 0.08). Full LMM results are provided in Supplementary Tables S4–S7.

**Figure 1.**
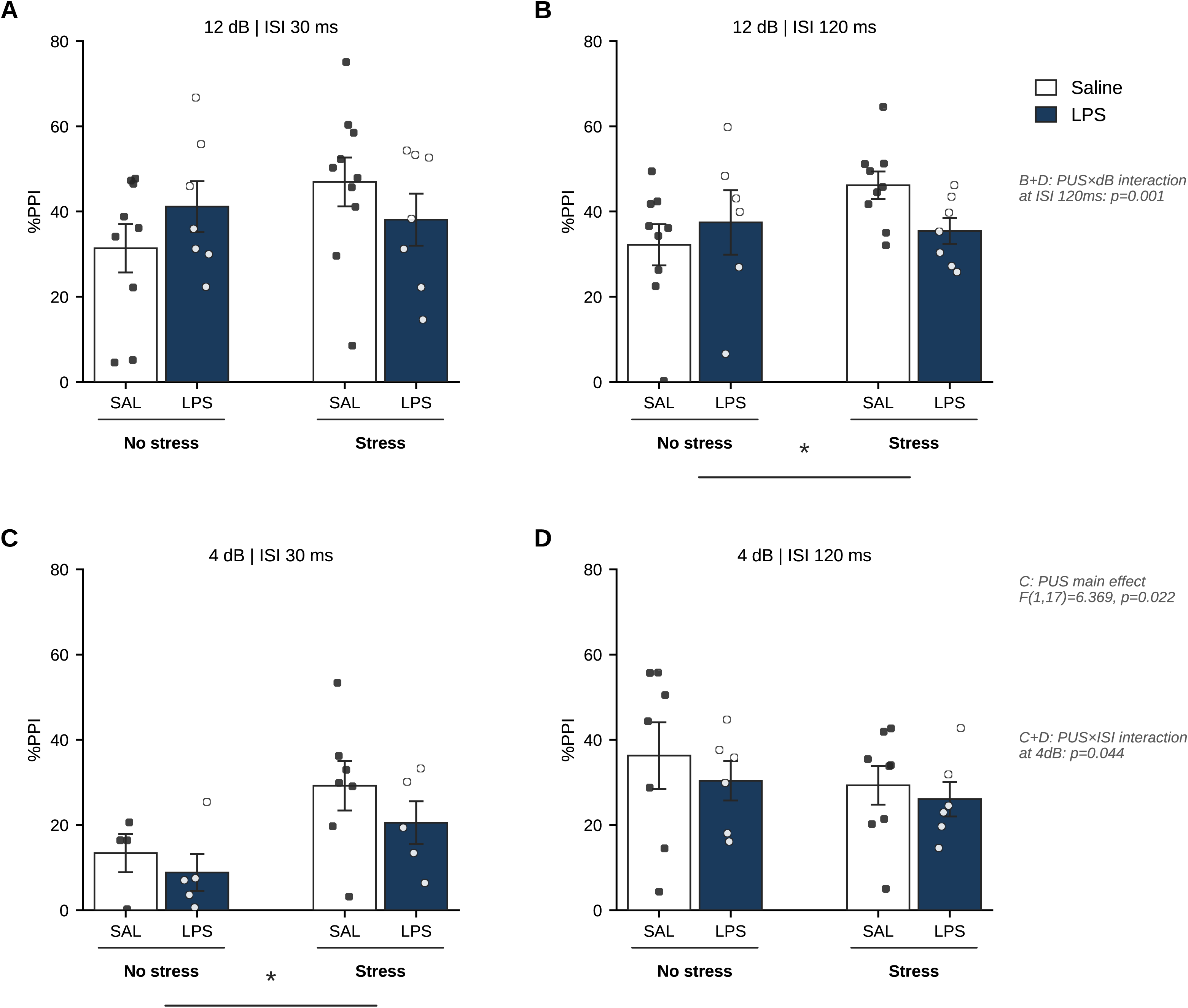
Prepulse inhibition (PPI) in adult male offspring of the maternal immune activation (MIA) and peripubertal stress (PUS) two-hit model. Each panel shows mean %PPI ± SEM for the four experimental groups (SAL/NS, LPS/NS, SAL/S, LPS/S) across conditions defined by prepulse intensity (12 dB or 4 dB above background noise) and inter-stimulus interval (ISI: 30 ms or 120 ms). Individual data points are superimposed. Negative PPI values were excluded prior to analysis. Linear mixed model analysis revealed a significant main effect of PUS at 12 dB (panels A and B; p = 0.027), a significant PUS × ISI interaction at 4 dB (panels C and D; p = 0.044), and a significant PUS × dB interaction at ISI 120 ms (panels B and D; p = 0.001). * p < 0.05 for PUS effect (horizontal line between No stress and Stress groups).

### 3.2 Glutamatergic system

RT-qPCR analysis revealed no significant effects of MIA or PUS on the expression of *Gria1* (GluA1; all p > 0.55), *Gria2* (GluA2; all p > 0.57), their ratio (all p > 0.40), or *Grm1* (mGluR1; all p > 0.31). Similarly, *Grin1* mRNA levels were not significantly affected by either MIA or PUS (all p > 0.26), nor were *Grin2a* (all p > 0.20) or *Grin2b* (MIA: F(1,31) = 0.094, p = 0.761; MIA × PUS: F(1,31) = 0.009, p = 0.923) expression levels individually altered. However, PUS significantly reduced the *Grin2a*/*Grin2b* mRNA ratio (F(1,30) = 12.875, p = 0.001; η^2^ =0.300) (Figure 2A), indicating a relative shift towards greater *Grin2b* expression in stressed animals irrespective of prenatal treatment (SAL/NS: 1.035 ± 0.042; SAL/S: 0.854 ± 0.030; LPS/NS: 1.022 ± 0.032; LPS/S: 0.919 ± 0.047). No significant MIA effect or MIA × PUS interaction was detected for this ratio (MIA: F(1,30) = 0.434, p = 0.515; interaction: F(1,30) = 0.981, p = 0.330). There was also a non-significant trend for PUS to increase *Grin2b* expression (F(1,31) = 3.486, p = 0.071), consistent with the ratio results (Supplementary Tables S10 and S12).

**Figure 2.**
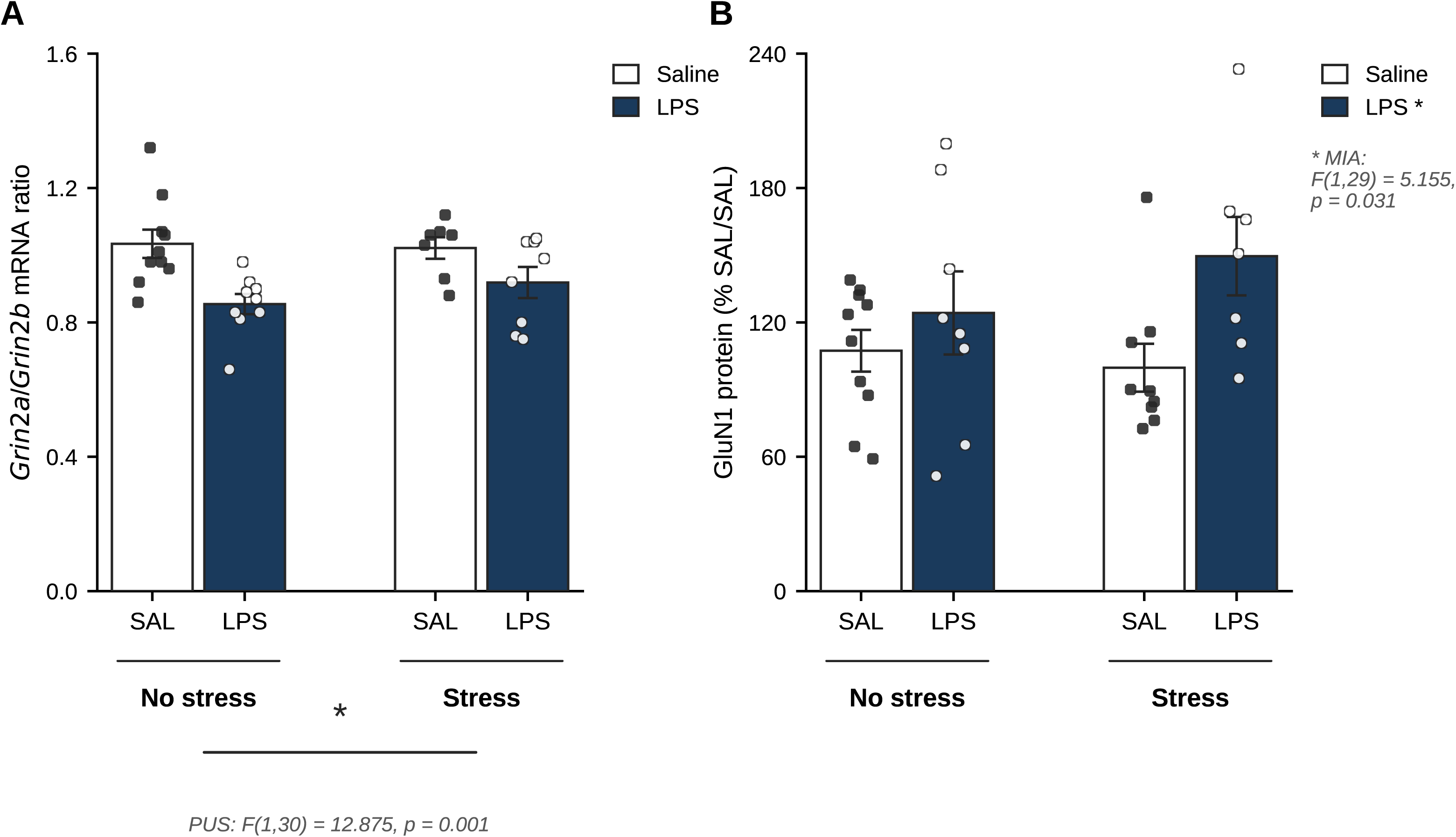
Effects of maternal immune activation (MIA) and peripubertal stress (PUS) on NMDA receptor markers in the adult rat dorsal hippocampus. (A) *Grin2a*/*Grin2b* mRNA ratio assessed by RT-qPCR, normalised to *Gapdh*. PUS significantly reduced the ratio, indicating a relative shift towards greater *Grin2b* expression. * p < 0.05 for PUS main effect (horizontal line between No stress and Stress groups). (B) GluN1 protein expression normalised to total protein load (Stain-Free). MIA significantly increased GluN1 protein levels (see Supplementary Table S14 and S15 for further details). * p < 0.05 for MIA main effect (indicated in legend). Data are mean ± SEM with individual data points superimposed. White bars: Saline; dark bars: LPS.

At the protein level, MIA significantly increased GluN1 expression in the dorsal hippocampus (Figure 2B; F(1,29) = 5.155, p = 0.031; η^2^ =0.151), with LPS-exposed animals showing higher GluN1 protein levels than saline controls regardless of postnatal treatment. Neither PUS nor the MIA × PUS interaction reached significance for GluN1 protein (PUS: F(1,29) = 0.474, p = 0.497; interaction: F(1,29) = 1.183, p = 0.286). Taken together, these findings suggest a dissociation between prenatal and peripubertal effects on hippocampal glutamatergic signalling: MIA selectively increased GluN1 protein without altering subunit composition at the mRNA level, while PUS altered the *Grin2a*/*Grin2b* ratio without affecting total receptor levels.

Neither MIA nor PUS significantly altered hippocampal glutamate, glutamine, or their ratio (all p > 0.39; Supplementary Tables S8-9).

### 3.3 GABA-A**γ**2 Subunit and GABA Levels

Neither MIA nor PUS significantly affected *Gabrg2* (GABA-Aγ2 subunit) mRNA expression in the dorsal hippocampus (MIA: F(1,30) = 1.705, p = 0.202; PUS: F(1,30) = 0.585, p = 0.450; MIA × PUS: F(1,30) = 0.021, p = 0.886). Similarly, hippocampal GABA concentrations measured by capillary electrophoresis were not significantly altered by either treatment (MIA: F(1,30) = 0.533, p = 0.471; PUS: F(1,30) = 0.996, p = 0.334; MIA × PUS: F(1,30) = 2.585, p = 0.118) (Supplementary tables S8, S9, S10 and S12).

### 3.4 Metabolites

MIA and PUS displayed interactive effects on taurine levels (Figure 3B) in the dorsal hippocampus (MIA × PUS: F(1,30) = 5.510, p = 0.026; η^2^_p_=0.155). Simple effects analyses with Bonferroni correction revealed that within the stressed animals, taurine was significantly elevated in LPS-exposed animals compared to saline controls (LPS/S: 107.3 ± 3.6 µM vs. SAL/S: 90.7 ± 5.5 µM), whereas no significant difference was found between non-stressed groups (SAL/NS: 99.8 ± 7.2 µM; LPS/NS: 90.7 ± 3.1 µM). Neither the main effect of MIA (F(1,30) = 0.472, p = 0.497) nor that of PUS (F(1,30) = 0.483, p = 0.492) reached significance.

**Figure 3.**
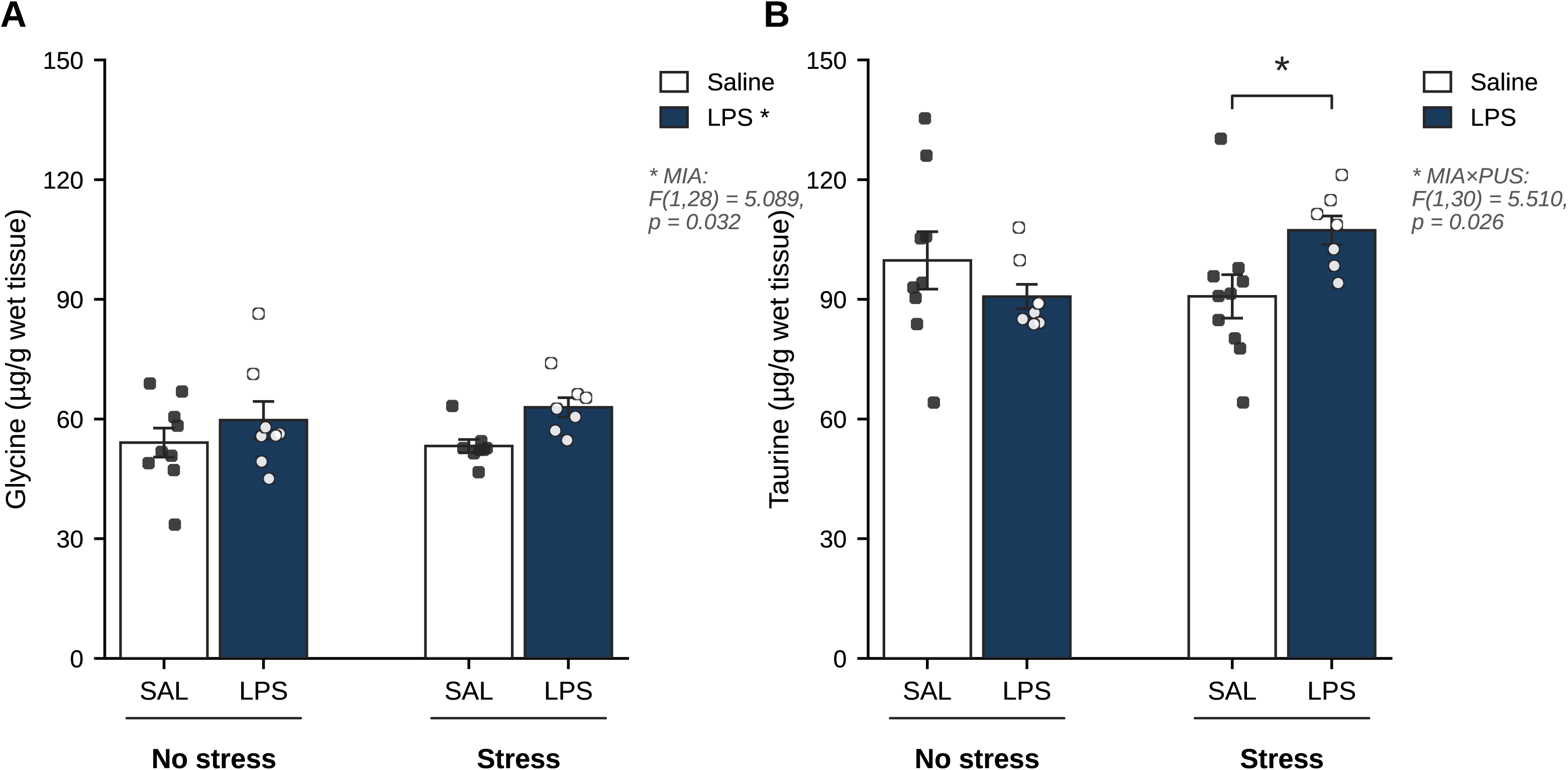
Effects of maternal immune activation (MIA) and peripubertal stress (PUS) on neuromodulatory amino acid levels in the adult rat dorsal hippocampus, measured by capillary electrophoresis with laser-induced fluorescence detection (CE-LIF). (A) Glycine concentrations. MIA significantly increased hippocampal glycine independently of PUS (see Supplementary Table S8 and S9 for further details). * p < 0.05 for MIA main effect (indicated in legend). (B) Taurine concentrations. A significant MIA × PUS interaction was detected; taurine was elevated in the LPS/S group compared to the SAL/S group within stressed animals. * p < 0.05 for simple effect within stressed animals (bracket). Data are mean ± SEM with individual data points superimposed. White bars: Saline; dark bars: LPS.

A significant main effect of MIA was found for glycine levels (Figure 3A) (F(1,28) = 5.089, p = 0.032; η^2^_p_0.154), with LPS-exposed animals showing higher hippocampal glycine concentrations than saline controls regardless of peripubertal stress exposure (SAL/NS: 54.1 ± 3.6 µM; SAL/stress: 53.2 ± 1.6 µM; LPS/NS: 59.7 ± 4.7 µM; LPS/S: 62.9 ± 2.4 µM). Neither PUS (F(1,28) = 0.116, p = 0.736) nor the MIA × PUS interaction (F(1,28) = 0.357, p = 0.555) reached significance. No significant effects were detected for L-glutamate, L-glutamine, L-serine, aspartate, or the L-glutamine/L-glutamate ratio (all p > 0.06; Table S8 and S9). A non-significant trend towards a MIA × PUS interaction was observed for aspartate (F(1,29) = 3.590, p = 0.068) and L-serine (F(1,30) = 3.208, p = 0.083), which may warrant further investigation.

### 3.5 Endocannabinoid system

PUS significantly increased *Mgll* (MAGL) mRNA expression (Figure 4B) in the dorsal hippocampus (F(1,30) = 4.494, p = 0.042; η2_p_=0.130), with stressed animals showing higher *Mgll* levels than non-stressed controls regardless of prenatal treatment (SAL/NS: 1.000 ± 0.096; SAL/S: 1.106 ± 0.113; LPS/NS: 0.813 ± 0.051; LPS/S: 1.113 ± 0.113). Neither MIA (F(1,30) = 0.890, p = 0.353) nor the MIA × PUS interaction (F(1,30) = 1.024, p = 0.320) was significant for *Mgll*.

**Figure 4.**
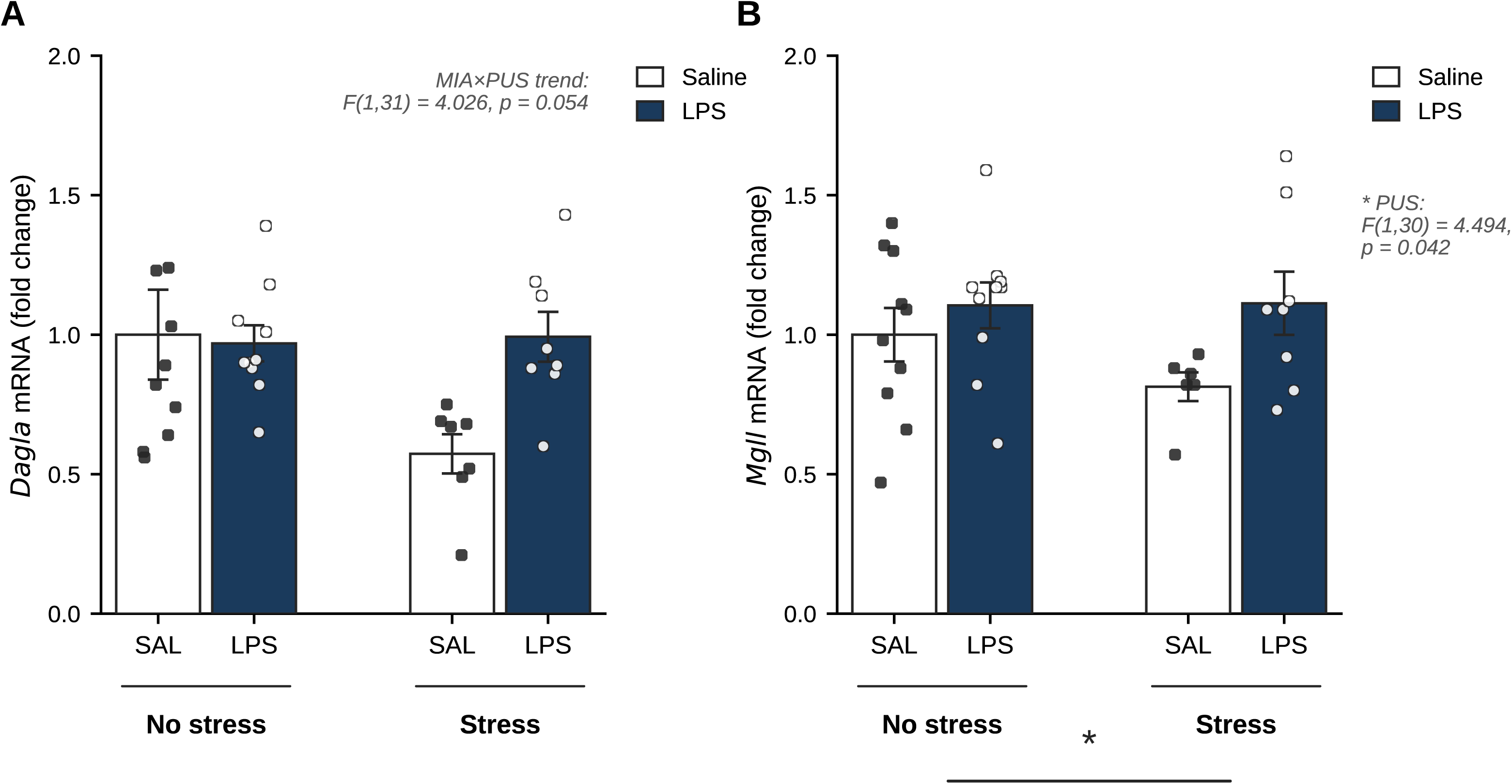
Effects of maternal immune activation (MIA) and peripubertal stress (PUS) on endocannabinoid system enzyme gene expression in the adult rat dorsal hippocampus, assessed by RT-qPCR and normalised to GAPDH. (A) *Dagla* (DAGLα) mRNA fold change. A near-significant MIA × PUS interaction trend was observed (F(1,31) = 4.026, p = 0.054), driven by reduced *Dagla* expression in LPS/NS animals relative to SAL/NS controls, an effect absent in stressed animals. (B) *Mgll* (MAGL) mRNA fold change. PUS significantly increased *Mgll* expression regardless of prenatal treatment (Supplementary Table S11 and S13). * p < 0.05 for PUS main effect (horizontal line between No stress and Stress groups). Data are mean ± SEM with individual data points superimposed. White bars: Saline; dark bars: LPS.

With regard to *Dagla* (DAGLα) expression, a near-significant MIA × PUS interaction was observed (Figure 4A; F(1,31) = 4.026, p = 0.054). Inspection of group means suggested that within the non-stressed animals, *Dagla* expression was lower in LPS-exposed animals than in saline controls (LPS/NS: 0.573 ± 0.070 vs. SAL/NS: 1.000 ± 0.161), an effect that was absent in stressed animals (LPS/S: 0.992 ± 0.090; SAL/S: 0.969 ± 0.065), though simple effects did not survive Bonferroni correction. Neither the main effect of MIA (F(1,31) = 3.198, p = 0.083) nor that of PUS (F(1,31) = 2.954, p = 0.096) reached significance independently.

No significant effects of MIA, PUS, or their interaction were detected for *Cnr1* (CB1 receptor; all p > 0.76), *Napepld* (NAPE-PLD; all p > 0.37), *Faah* (FAAH; all p > 0.24), the *Mgll*/*Dagla* ratio (all p > 0.39), or the *Napepld*/*Faah* ratio (all p > 0.34; Table S11 and S13).

### 3.6 nNOS

Real time qPCR showed no significant effects of MIA ((F (1,28) = 1.746; p=0.197)), PUS ((F (1,28) = 2.223; p=0.147) or their interaction ((F (1,29) = 0.148; p=0.704)). Concordantly, western blot analysis revealed no significant effects of MIA, PUS, or their interaction on nNOS protein expression in the adult rat dorsal hippocampus (MIA: F(1,29) = 0.352, p = 0.557; PUS: F(1,29) = 0.470, p = 0.499; MIA × PUS: F(1,29) = 0.148, p = 0.704; Table S14 and S15).

Table 1 provides a summary of the key findings of the study.

**Table 1.** Summary of significant and trend-level effects for all variables measured in the adult rat dorsal hippocampus.

| System | Variable | Level | MIA effect | PUS effect | MIA x PUS | Key finding |
| --- | --- | --- | --- | --- | --- | --- |
| Sensorimotor gating | *PPI4_30* | Behaviour |  | * |  | PUS increases PPI at 4 dB / 30 ms ISI |
|  | *PPI at 12 dB (a)* | Behaviour |  | * |  | PUS increases PPI at 12 dB (LMM) |
| Glutamatergic system | *Grin2a/Grin2b ratio* | mRNA |  | * |  | PUS reduces ratio: shift towards GluN2B (immature NMDAR) |
|  | *GluN1* | Protein | * |  |  | MIA increases GluN1 protein expression |
| GABAergic system | *Gabrg2, GABA* | mRNA / Metabolite |  |  |  | No significant effects |
| Neuromodulatory amino acids | *Glycine* | Metabolite | * |  |  | MIA increases glycine independently of PUS |
|  | *Taurine* | Metabolite |  |  | * | LPS/S > SAL/S within stressed animals |
| Endocannabinoid system | *Mgll (MAGL)* | mRNA |  | * |  | PUS increases MAGL expression |
|  | *Dagla (DAGLa)* | mRNA |  |  | (+) | Trend: MIA reduces DAGLa in non-stressed animals only |
|  | *Cnr1, Napepld, Faah* | mRNA |  |  |  | No significant effects |
| Nitroergic system | *nNOS* | Protein |  |  |  | No significant effects |

\* Significant ( $p < 0.05$ ) (+) trend ( $p < 0.10$ ) (a) PPI at 12 dB by linear mixed model (LMM); all others by two-way ANOVA (MIA x PUS). MIA: maternal immune activation (LPS, GD15-16); PUS: peripubertal stress (PND28-38); LPS/S: LPS stressed; SAL/S: saline stressed.

## 4. Discussion

The present study examined the long-term neurochemical consequences of MIA and PUS, alone and in combination, on multiple signalling systems in the adult rat dorsal hippocampus. Our findings reveal a pattern of complementary but mechanistically distinct disruptions: MIA selectively increased GluN1 protein expression and elevated hippocampal glycine levels, while PUS reduced the *Grin2a*/*Grin2b* mRNA ratio and upregulated *Mgll* expression. A significant MIA × PUS interaction was detected for taurine, with elevation specific to the double-hit condition. Notably, neither GABAergic markers nor nNOS protein were altered by either insult. In addition, while no deficits in PPI were observed due to MIA, PUS increased PPUI at high prepulse intensities. We suggest that these results may be consistent with a model in which early developmental insults produce a state of latent hippocampal hyperexcitability through glutamatergic upregulation and incomplete synaptic maturation, without adequate inhibitory or endocannabinoid compensation, and with neuromodulatory amino acids reflecting late-stage homeostatic attempts to restore E/I balance.

### 4.1 Prepulse inhibition

Linear mixed model analysis revealed that PUS consistently facilitated PPI at high prepulse intensities (12 dB above background, both ISIs; p = 0.027), while its effects at low intensities (4 dB) were ISI-dependent: PUS increased PPI at 30 ms onset asynchrony but reduced it at 120 ms, a pattern that produces mutual cancellation when conditions are averaged. A PUS × dB interaction at ISI 120 ms further confirmed that the direction of stress effects on sensorimotor gating depends on stimulus parameters. No significant effects of MIA were detected, though a trend towards a MIA × PUS interaction at 12 dB may warrant further investigation with larger samples.

The consistent finding is therefore that PUS facilitated rather than impaired sensorimotor gating — a result that does not follow the PPI deficit pattern sometimes expected in two-hit neurodevelopmental models but is fully consistent with the broader literature in which PPI outcomes are highly variable across model configurations. PPI deficits have been demonstrated in two-hit models combining MIA with peripubertal stress — specifically in protocols using poly I:C combined with heterotypic peripubertal stressors similar to those used here (Giovanoli et al., 2013; Deslauriers et al., 2013) — but these effects are not universal across two-hit designs. In particular, models combining prenatal MIA with adolescent cannabinoid exposure have consistently failed to produce PPI deficits (Martin-Cuevas et al., 2023; Moreno-Fernández et al., 2024, 2025), and our own previous work using the same MIA + PUS protocol in rats did not detect PPI alterations (Capellán et al., 2023). This pattern suggests that PPI disruption in two-hit models is not a generic consequence of combining two developmental insults, but depends critically on the nature of each hit, its timing, the species and strain used, and the acoustic stimulus parameters employed (Khan and Powell, 2017). Although intense or repeated stress consistently impairs PPI through dopaminergic mechanisms, moderate stress levels — particularly during sensitive developmental windows — have been shown to enhance sensorimotor gating, following an inverted-U dose-response pattern (Pujante-Gil et al., 2021). The heterotypic peripubertal protocol used here may therefore have produced a facilitating rather than impairing effect at specific stimulus intensities, consistent with the hypothesis that stress inoculation during sensitive developmental periods promotes more resilient behavioural phenotypes (Pujante-Gil et al., 2021). Importantly, the absence of PPI deficits does not preclude meaningful neurochemical dysregulation, as sensorimotor gating and molecular markers of E/I balance can dissociate across measures and are not necessarily co-expressed following the same developmental insults. The analysis of prepulse facilitation rates further supports this interpretation: the proportion of negative PPI values was primarily determined by stimulus parameters rather than by treatment, ruling out a differential effect of MIA or PUS on prepulse facilitation per se.

### 4.2 Glutamatergic system

The most consistent neurochemical alterations observed in this study were within the glutamatergic system, and they suggest a dissociation between the effects of prenatal and peripubertal insults on distinct aspects of NMDA receptor regulation. MIA significantly increased GluN1 protein (Figure 2B; Supplementary Table S14-15) expression in the dorsal hippocampus, without a corresponding change in *Grin1* mRNA, pointing to post-transcriptional or translational upregulation of this obligatory NMDA receptor subunit. This finding echoes previous reports in which increased GluN1 protein was detected following MIA despite unaltered mRNA levels (Rahman et al., 2017), and is consistent with the notion that MIA triggers lasting changes in receptor abundance through mechanisms that operate downstream of transcription. Increased GluN1 expression may reflect a compensatory upregulation in response to LPS-induced NMDA receptor hypofunction — a well-documented consequence of prenatal LPS exposure at the synaptic level (Burt et al., 2013) — that ultimately results in greater receptor availability in adulthood. Glutamatergic dysfunction is a core feature of neurodevelopmental disorders, with growing evidence linking NMDA receptor alterations to the cognitive deficits, impaired synaptic plasticity, and disrupted E/I balance observed in these conditions (Moghaddam and Javitt, 2012; Paoletti et al., 2013).

Independently of MIA, PUS significantly reduced the *Grin2a*/*Grin2b* mRNA ratio (Figure 2A; Supplementary Tables S10 and S12) in the dorsal hippocampus, without altering the individual expression of either subunit to a statistically significant degree. This reduction reflects a relative shift towards greater *Grin2b* expression, indicative of an incomplete or reversed GluN2B-to-GluN2A developmental switch. This maturational transition, which normally occurs during late postnatal and adolescent development, is essential for the stabilisation of excitatory synapses, as GluN2A-containing receptors confer faster kinetics, lower calcium permeability, and more precise synaptic plasticity thresholds than their GluN2B-containing counterparts (Paoletti et al., 2013; Yashiro and Philpot, 2008). Disruption of this switch has been linked to stress-induced interference with REST-mediated transcriptional control (Rodenas-Ruano et al., 2012), and a persistent GluN2B-rich receptor composition has been associated with increased excitability, impaired learning precision, and heightened vulnerability to excitotoxic insults (Yashiro and Philpot, 2008; Ladagu et al., 2023). Importantly, neither MIA nor the MIA × PUS interaction reached significance for this ratio, indicating that the effect of PUS on subunit maturation is independent of prenatal immune priming under the conditions tested. No significant effects were detected for AMPA receptor subunits (*Gria1*, *Gria2*) or the metabotropic receptor *Grm1*. Reductions in GluA1 protein levels have been reported in the hippocampus of poly I:C-exposed animals (Shao et al., 2023), though the literature on AMPA subunit changes following MIA is inconsistent and effects may be confined to specific hippocampal subregions not resolvable with whole-tissue analysis. Taken together, MIA and PUS appear to act on complementary yet distinct aspects of hippocampal glutamatergic signalling — receptor abundance and subunit maturation, respectively.

### 4.3 GABAergic system

Neither GABA-Aγ2 subunit mRNA expression nor hippocampal GABA concentrations were significantly altered by MIA, PUS, or their interaction. These null results are noteworthy in the context of the glutamatergic alterations described above, as they suggest that the hippocampal inhibitory system does not mount a detectable compensatory response to the increased excitatory tone produced by these insults — at least not at the level of the γ2 subunit or total GABA content. The γ2 subunit is critical for both GABA-A receptor assembly and its proper synaptic localisation, processes fundamental to inhibitory synapse function and plasticity (Schweizer et al., 2003; Chua and Chebib, 2017). Its stability in the present study could also reflect the specificity of two-hit effects to discrete hippocampal subregions — particularly CA3 and the dentate gyrus — that are not resolvable in whole-tissue analyses. Indeed, previous work has shown that reductions in GABAergic markers such as parvalbumin and reelin following combined MIA and PUS are confined to the ventral dentate gyrus (Giovanoli et al., 2014), and a systematic review of 102 MIA studies confirmed that GABAergic alterations are characteristically region-, timing-, and sex-specific (Gillespie et al., 2024). An alternative explanation is that any GABAergic changes induced by these insults are transient, occurring during early postnatal development or adolescence and normalising by adulthood. This temporal pattern has been described for several GABAergic markers in MIA models (Gillespie et al., 2024) and would be consistent with the idea that the hippocampus retains some capacity for inhibitory compensation in the short term, even if this proves insufficient over the longer developmental trajectory. The lack of inhibitory adaptation in the face of glutamatergic upregulation nonetheless supports the concept of a failed homeostatic E/I balancing response, in which the excitatory and inhibitory systems are decoupled rather than co-regulated (Sohal and Rubenstein, 2019; Uliana et al., 2024). While clinical studies report a global reduction in GABA across brain regions in schizophrenia spectrum disorders (Egerton et al., 2017), our findings do not support a persistent hippocampal GABA deficit under the conditions tested, a discrepancy that could reflect methodological differences between in vivo MRS measurements and post-mortem tissue homogenate analyses, or the distinct neurobiological consequences of the specific two-hit protocol used here.

### 4.4 Neuromodulatory amino acids

Two neuromodulatory amino acids were significantly altered: taurine, which showed a MIA × PUS interaction, and glycine, which was elevated by MIA independently of PUS. Both changes are consistent with compensatory responses to the heightened excitatory environment produced by these insults, though they differ in their specificity and putative neurochemical mediators.

Taurine was elevated selectively in the double-hit (LPS/S) group compared to SAL/S animals, with no significant difference between non-stressed groups. Taurine is an inhibitory neuromodulator with multiple neuroprotective functions: it regulates intracellular calcium homeostasis, limits oxidative stress, and attenuates excitotoxic damage, in part through its actions on GABA-A and glycine receptors (Wu and Prentice, 2010). Its selective elevation in the double-hit condition suggests a context-dependent regulatory response in which peripubertal stress, when superimposed on a prenatal immune-primed hippocampus, triggers adaptive taurinergic signalling aimed at counterbalancing the cumulative excitatory burden. This interpretation is consistent with the broader literature showing bidirectional changes in taurine depending on the nature and timing of the environmental insult (El-Maraghi et al., 2018; Zhu et al., 2022). It is also noteworthy that the observed taurine elevation is specific to the combination of MIA and PUS — neither insult alone produced a significant change — which mirrors the synergistic pattern of the two-hit hypothesis and suggests that taurine upregulation may be a marker of cumulative neurochemical stress rather than a response to either insult in isolation.

Hippocampal glycine was significantly elevated by MIA regardless of subsequent peripubertal stress, indicating a long-lasting effect of maternal immune activation on glycinergic signalling. Glycine serves dual roles in the CNS: as an inhibitory neurotransmitter via strychnine-sensitive glycine receptors (GlyR), and as an obligatory co-agonist at the NMDA receptor glycine-binding site, where it facilitates receptor activation (Balu, 2016). Its elevation following MIA may therefore represent an endogenous compensatory mechanism aimed at potentiating NMDA receptor function in the context of LPS-induced receptor hypofunction (Burt et al., 2013). This interpretation is supported by clinical observations linking glycine modulation to NMDA receptor function in schizophrenia, where glycine-site agonists and GlyT1 transport inhibitors have been explored as adjunctive treatments (Cavalcante et al., 2026). However, because glycine preferentially potentiates GluN2B-containing receptors — which display high glycine affinity — its elevation in animals that also show a PUS-induced relative increase in GluN2B expression could contribute to a maladaptive feedback loop that further amplifies excitatory drive. Although the effects of MIA (elevated GluN1, elevated glycine) and PUS (reduced *Grin2a*/*Grin2b* ratio) were largely independent, their co-occurrence in the double-hit condition, perhaps at a different timepoint from the one assessed here, may nonetheless produce a hippocampal environment with both greater NMDA receptor density and a more excitable subunit composition — a combination that could contribute to latent synaptic vulnerability, and whose cumulative impact is further reflected in the taurine elevation specific to the LPS/S group. Elevated hippocampal glycine has been observed in other rodent models of maternal immune activation (Di Maio et al., 2025), supporting the specificity of this finding to the MIA component of the two-hit protocol.

### 4.5 Endocannabinoid system

Peripubertal stress significantly increased *Mgll* (MAGL) expression in the dorsal hippocampus, without affecting *Dagla* (DAGLα), *Cnr1* (CB1), *Napepld* (NAPE-PLD), or *Faah* expression. A near-significant MIA × PUS interaction was observed for *Dagla* (p = 0.054), with reduced expression selectively in the LPS/NS group compared to SAL/NS, an effect that was absent in stressed animals. MAGL is the primary enzyme responsible for the degradation of 2-arachidonoylglycerol (2-AG), the main endocannabinoid ligand at CB1 receptors, while DAGLα is its principal biosynthetic enzyme. Upregulation of MAGL following PUS would be expected to reduce 2-AG availability, thereby diminishing CB1-mediated retrograde inhibition of both glutamatergic and GABAergic synapses. This finding is consistent with prior reports that chronic unpredictable stress upregulates MAGL in the hippocampus (Fang and Wang, 2018) which may lead to reduced endocannabinoid tone and impaired stress resilience. MAGL inhibition has been shown to reverse stress-induced hippocampal LTP impairment and to normalise adult neurogenesis deficits (Zhang et al., 2015), further underscoring the functional importance of this enzyme in hippocampal plasticity under stress.

The endocannabinoid system is increasingly implicated in the pathophysiology of neurodevelopmental disorders, given its key role in modulating synaptic plasticity, stress responsivity, and E/I balance (Navarrete et al., 2020; Santoni and Pistis, 2025). Reductions in CB1 receptor expression have been reported in post-mortem schizophrenia brain tissue and in animal models of MIA (Bloch Priel et al., 2023; Santoni and Pistis, 2025), and altered 2-AG signalling has been proposed as a compensatory response to neuroinflammation following MIA (Guo et al., 2018; Santoni and Pistis, 2025). The absence of CB1 changes in the present study is consistent with findings from other two-hit models in which receptor expression remains stable despite altered enzymatic regulation (Santoni and Pistis, 2025), and suggests that the ECS alterations observed here may be specific to 2-AG metabolism rather than to receptor-level remodelling. The trend towards reduced *Dagla* in non-stressed LPS animals, if confirmed with larger samples, would suggest that MIA may suppress 2-AG synthesis in the absence of subsequent stress — an effect that peripubertal stress appears to normalise. This pattern would be compatible with a two-hit model in which the first hit (MIA) reduces anandabinoid tone through impaired synthesis, and the second hit (PUS) superimposes accelerated degradation via MAGL, resulting in a net reduction of 2-AG availability that reduces the capacity for retrograde inhibition and may thereby contribute to hippocampal hyperexcitability.

### 4.6 Neuronal nitric oxide synthase

No significant effects of MIA, PUS, or their interaction were detected for nNOS protein (Supplementary Table S14 and S15) expression in the adult dorsal hippocampus. This null result is informative rather than simply negative, as it is consistent with emerging evidence that nNOS alterations following MIA are developmentally transient and highly region-specific. Zhang et al. (2017) reported increased nNOS immunoreactivity in the CA3 region and dentate gyrus of neonatal rat offspring at postnatal day 2 following poly I:C MIA, but not in the CA1 region; and Tellez-Merlo et al. (2019), using a prenatal LPS protocol closely comparable to the present study, found elevated NO levels in the prefrontal cortex, brainstem, and amygdala, but not in the dorsal hippocampus of post-pubertal offspring. Taken together, these findings suggest that any MIA-induced perturbation of hippocampal nNOS normalises by adulthood, or is restricted to subregions and cellular compartments not detectable in whole-tissue protein analyses. nNOS is expressed predominantly in a subpopulation of GABAergic interneurons in the hippocampus (Tricoire and Vitalis, 2012), and its activity is tightly regulated by NMDA receptor-dependent calcium influx (Hardingham et al., 2013). It is therefore possible that nNOS function — rather than expression — is altered in our model, a hypothesis that could be addressed by measuring NO metabolites, nNOS phosphorylation state, or examining its co-expression with specific interneuron markers. Despite being encoded by NOS1, a recognised genetic risk factor for schizophrenia-spectrum disorders (Nasyrova et al., 2015; Freudenberg et al., 2015), and having been implicated in sensorimotor gating deficits (Rovný et al., 2018), our data do not support a role for nNOS protein dysregulation in the adult dorsal hippocampus as a consequence of this specific two-hit protocol.

### 4.7 Limitations and future directions

Several limitations of the present study should be acknowledged. First, the analysis was performed on whole dorsal hippocampal homogenates, which precludes the detection of subregion-specific effects in CA1, CA3, or the dentate gyrus — regions that may respond differentially to MIA and PUS, as suggested by the literature on GABAergic and nNOS markers. Second, only male offspring were included, limiting the generalisability of the findings given the well-documented sex-dependent effects of MIA on glutamatergic and GABAergic signalling (Rahman et al., 2017; Gillespie et al., 2024). Third, the moderate sample sizes per group (n = 7–10) may have reduced statistical power for detecting smaller effect sizes, particularly for the near-significant *Dagla* interaction. Fourth, the present study assessed gene and protein expression as proxies for receptor and enzyme activity, but did not directly measure synaptic function, endocannabinoid levels, or amino acid release, which would be required to confirm the functional consequences of the molecular alterations observed. Future studies should examine these parameters using electrophysiological and in vivo neurochemical approaches, include female offspring, and extend the analysis to hippocampal subregions and to other brain structures such as the prefrontal cortex and ventral hippocampus, which are closely interconnected with the dorsal hippocampus and may show complementary or divergent patterns of dysregulation.

### 4.8 Conclusions

In summary, maternal immune activation and peripubertal stress exert distinct and partially synergistic effects on hippocampal neurochemistry in adult male rats. MIA selectively upregulates GluN1 protein and hippocampal glycine — both consistent with a compensatory response to prenatal NMDA receptor hypofunction — while PUS disrupts the maturation of NMDA receptor subunit composition and reduces endocannabinoid degradative capacity. Neither insult alone produces detectable changes in GABAergic markers or nNOS, pointing to the specificity of these systems to particular insult combinations, subregions, or developmental windows. The significant MIA × PUS interaction for taurine, with elevation specific to the double-hit condition, provides the clearest evidence of a synergistic neurochemical phenotype and identifies taurine as a potential marker of cumulative excitatory stress. Taken together, these findings support a model of latent hippocampal hyperexcitability in which complementary glutamatergic adaptations converge without adequate inhibitory counterbalance. These findings add to the existing evidence implicating these systems in neurodevelopmental vulnerability, and highlight their interactive dysregulation in the dorsal hippocampus as a specific target for future mechanistic investigation.

## Acknowledgments

We would like to acknowledge Rosa Ferrado for the excellent technical assistance.

## References

Balu, D.T., 2016. The NMDA receptor and schizophrenia: from pathophysiology to treatment. Adv. Pharmacol. 76, 351–382. 10.1016/bs.apha.2016.01.006

Bannerman, D.M., Rawlins, J.N., McHugh, S.B., Deacon, R.M., Yee, B.K., Bast, T., Zhang, W.N., Pothuizen, H.H., Feldon, J., 2004. Regional dissociations within the hippocampus — memory and anxiety. Neurosci. Biobehav. Rev. 28, 273–283. 10.1016/j.neubiorev.2004.03.004

Batinić, B., Santrač, A., Divović, B., Timić, T., Stanković, T., Obradović, ALj., Joksimović, S., Savić, MM., 2016. Lipopolysaccharide exposure during late embryogenesis results in diminished locomotor activity and amphetamine response in females and spatial cognition impairment in males in adult, but not adolescent rat offspring. Behav Brain Res. Feb 15;299:72–80. doi: 10.1016/j.bbr.2015.11.025.

Bloch Priel, S., Yitzhaky, A., Gurwitz, D., Hertzberg, L., 2023. Cannabinoid receptor gene CNR1 is downregulated in subcortical brain samples and upregulated in blood samples of individuals with schizophrenia: a participant data systematic meta-analysis. Eur. J. Neurosci. 58, 3540–3554. 10.1111/ejn.16122

Boley, A.M., Perez, S.M., Lodge, D.J., 2014. A fundamental role for hippocampal parvalbumin in the dopamine hyperfunction associated with schizophrenia. Schizophr. Res. 157, 238–243. 10.1016/j.schres.2014.05.005

Borsini, A., Giacobbe, J., Mandal, G., Boldrini, M., 2023. Acute and long-term effects of adolescence stress exposure on rodent adult hippocampal neurogenesis, cognition, and behaviour. Mol. Psychiatry 28, 4124–4137. 10.1038/s41380-023-02229-2

Braff, D.L., Geyer, M.A., Swerdlow, N.R., 2001. Human studies of prepulse inhibition of startle: normal subjects, patient groups, and pharmacological studies. Psychopharmacology 156, 234–258. 10.1007/s002130100810

Burt, M.A., Tse, Y.C., Boksa, P., Wong, T.P., 2013. Prenatal immune activation interacts with stress and corticosterone exposure later in life to modulate N-methyl-D-aspartate receptor synaptic function and plasticity. Int. J. Neuropsychopharmacol. 16, 1835–1848. 10.1017/S1461145713000229

Capellán, R., Moreno-Fernández, M., Orihuel, J., Roura-Martínez, D., Ucha, M., Ambrosio, E., Higuera-Matas, A., 2022. Ex vivo ¹H-MRS brain metabolic profiling in a two-hit model of neurodevelopmental disorders: prenatal immune activation and peripubertal stress. Schizophr. Res. 243, 232–240. 10.1016/j.schres.2019.11.007

Capellán, R., Orihuel, J., Marcos, A., Ucha, M., Moreno-Fernández, M., Casquero-Veiga, M., Soto-Montenegro, M.L., Desco, M., Oteo-Vives, M., Ibáñez-Moragues, M., Magro-Calvo, N., Morcillo, M.Á., Ambrosio, E., Higuera-Matas, A., 2023. Interaction between maternal immune activation and peripubertal stress in rats: impact on cocaine addiction-like behaviour, morphofunctional brain parameters and striatal transcriptome. Transl. Psychiatry 13, 84. 10.1038/s41398-023-02378-6

Cavalcante, D.P., Carvalho, G.A., Nunes, A.Í.S., Quintanilha, A.R., Nascimento, L.R.C., Caixeta, L., Ulrich, H., Gomez, R.S., Pinto, M.C.X., 2026. GlyT1 (SLC6A9) inhibition in neurological and psychiatric disorders. Naunyn Schmiedebergs Arch. Pharmacol. 10.1007/s00210-026-05276-y

Chua, H.C., Chebib, M., 2017. GABAA receptors and the diversity in their structure and pharmacology. Adv. Pharmacol. 79, 1–34. 10.1016/bs.apha.2017.03.003

Deslauriers, J., Larouche, A., Sarret, P., Grignon, S., 2013. Combination of prenatal immune challenge and restraint stress affects prepulse inhibition and dopaminergic/GABAergic markers. Prog. Neuropsychopharmacol. Biol. Psychiatry 45, 156–164. 10.1016/j.pnpbp.2013.05.006

Di Maio, A., Yahyavi, I., Buzzelli, V., Motta, Z., Ascone, F., Putignani, L., Usiello, A., Pollegioni, L., Trezza, V., Errico, F., 2025. Prenatal exposure to lipopolysaccharide or valproate leads to abnormal accumulation of the NMDA receptor agonist D-aspartate in the adolescent rat brain. J. Neurochem. 169, e70095. 10.1111/jnc.70095

Egerton, A., Modinos, G., Ferrera, D., McGuire, P., 2017. Neuroimaging studies of GABA in schizophrenia: a systematic review with meta-analysis. Transl. Psychiatry 7, e1147. 10.1038/tp.2017.124

El-Maraghi, E.F., Abdel-Fattah, K.I., Soliman, S.M., El-Sayed, W.M., 2018. Taurine provides a time-dependent amelioration of the brain damage induced by γ-irradiation in rats. J. Hazard. Mater. 359, 40–46. 10.1016/j.jhazmat.2018.07.005

Escobar, M., Crouzin, N., Cavalier, M., Quentin, J., Roussel, J., Lanté, F., Batista-Novais, A.R., Cohen-Solal, C., De Jesus Ferreira, M.C., Guiramand, J., Barbanel, G., Vignes, M., 2011. Early, time-dependent disturbances of hippocampal synaptic transmission and plasticity after in utero immune challenge. Biol. Psychiatry 70, 992–999. 10.1016/j.biopsych.2011.01.009

Fang, G., Wang, Y. 2018. Effects of rTMS on Hippocampal Endocannabinoids and Depressive-like Behaviors in Adolescent Rats. Neurochem Res. Sep;43(9):1756–1765. doi: 10.1007/s11064-018-2591-y.

Fanselow, M.S., Dong, H.W., 2010. Are the dorsal and ventral hippocampus functionally distinct structures? Neuron 65, 7–19. 10.1016/j.neuron.2009.11.031

Freudenberg, F., Alttoa, A., Reif, A., 2015. Neuronal nitric oxide synthase (NOS1) and its adaptor, NOS1AP, as genetic risk factors for psychiatric disorders. Genes Brain Behav. 14, 46–63. 10.1111/gbb.12193

Gillespie, B., Panthi, S., Sundram, S., Hill, R.A., 2024. The impact of maternal immune activation on GABAergic interneuron development: a systematic review of rodent studies and their translational implications. Neurosci. Biobehav. Rev. 156, 105488. 10.1016/j.neubiorev.2023.105488

Giovanoli, S., Engler, H., Engler, A., Richetto, J., Voget, M., Willi, R., Winter, C., Riva, M.A., Mortensen, P.B., Feldon, J., Schedlowski, M., Meyer, U., 2013. Stress in puberty unmasks latent neuropathological consequences of prenatal immune activation in mice. Science 339, 1095–1099. 10.1126/science.1228261

Giovanoli, S., Weber, L., Meyer, U., 2014. Single and combined effects of prenatal immune activation and peripubertal stress on parvalbumin and reelin expression in the hippocampal formation. Brain Behav. Immun. 40, 48–54. 10.1016/j.bbi.2014.04.005

Guo, Z., Tse, Y.C., Zhang, Y., Sun, Q., Vecchiarelli, H.A., Aukema, R., Hill, M.N., Wong, T.P., Boksa, P., 2018. Prenatal immune activation potentiates endocannabinoid-related plasticity of inhibitory synapses in the hippocampus of adolescent rat offspring. Eur. Neuropsychopharmacol. 28, 1405–1417. 10.1016/j.euroneuro.2018.09.003

Hardingham, N., Dachtler, J., Fox, K., 2013. The role of nitric oxide in pre-synaptic plasticity and homeostasis. Front. Cell. Neurosci. 7, 190. 10.3389/fncel.2013.00190

Heckers, S., Konradi, C., 2015. GABAergic mechanisms of hippocampal hyperactivity in schizophrenia. Schizophr. Res. 167, 4–11. 10.1016/j.schres.2014.09.041

Ito, H.T., Smith, S.E.P., Hsiao, E., Patterson, P.H., 2010. Maternal immune activation alters nonspatial information processing in the hippocampus of the adult offspring. Brain Behav. Immun. 24, 930–941. 10.1016/j.bbi.2010.03.004

Khan, A., Powell, S.B., 2017. Sensorimotor gating deficits in two-hit models of schizophrenia risk factors. Schizophr. Res. 198, 68–83. 10.1016/j.schres.2017.10.009

Kiemes, A., Davies, C., Kempton, M.J., Lukow, P.B., Bennallick, C., Stone, J.M., Modinos, G., 2021. GABA, glutamate and neural activity: a systematic review with meta-analysis of multimodal 1H-MRS-fMRI studies. Front. Psychiatry 12, 644315. 10.3389/fpsyt.2021.644315

Konradi, C., Yang, C.K., Zimmerman, E.I., Lohmann, K.M., Gresch, P., Pantazopoulos, H., Berretta, S., Heckers, S., 2011. Hippocampal interneurons are abnormal in schizophrenia. Schizophr. Res. 131, 165–173. 10.1016/j.schres.2011.06.007

Ladagu, A.D., Olopade, F.E., Adejare, A., Olopade, J.O., 2023. GluN2A and GluN2B N-methyl-D-aspartate receptor subunits: their roles and therapeutic antagonists in neurological diseases. Pharmaceuticals 16, 1535. 10.3390/ph16111535

Lewis, D.A., Curley, A.A., Glausier, J.R., Volk, D.W., 2012. Cortical parvalbumin interneurons and cognitive dysfunction in schizophrenia. Trends Neurosci. 35, 57–67. 10.1016/j.tins.2011.10.004

Lisman, J.E., Coyle, J.T., Green, R.W., Javitt, D.C., Benes, F.M., Heckers, S., Grace, A.A., 2008. Circuit-based framework for understanding neurotransmitter and risk gene interactions in schizophrenia. Trends Neurosci. 31, 234–242. 10.1016/j.tins.2008.02.005

Lorenzo, M.P., Villaseñor, A., Ramamoorthy, A., Garcia, A., 2013. Optimization and validation of a CE-LIF method for amino acids determination in human plasma. Application to bipolar disorder study. Electrophoresis 34, 1701–1709. 10.1002/elps.201200632

Martín-Cuevas, C., Ramos-Herrero, V.D., Crespo-Facorro, B., Sánchez-Hidalgo, A.C., 2023. Prenatal risk factors and postnatal cannabis exposure: assessing dual models of schizophrenia-like rodents. Neurosci. Biobehav. Rev. 154, 105409. 10.1016/j.neubiorev.2023.105409

McCormick, C.M., Thomas, C.M., Sheridan, C.S., Nixon, F., Flynn, J.A., Mathews, I.Z., 2012. Social instability stress in adolescent male rats alters hippocampal neurogenesis and produces deficits in spatial location memory in adulthood. Hippocampus 22, 1300–1312. 10.1002/hipo.20966

Meyer, U., 2019. Neurodevelopmental resilience and susceptibility to maternal immune activation. Trends Neurosci. 42, 793–806. 10.1016/j.tins.2019.08.001

Moghaddam, B., Javitt, D., 2012. From revolution to evolution: the glutamate hypothesis of schizophrenia and its implication for treatment. Neuropsychopharmacology 37, 4–15. 10.1038/npp.2011.181

Moreno-Fernández, M., Ucha, M., Reis-de-Paiva, R., Marcos, A., Ambrosio, E., Higuera-Matas, A., 2024. Lack of interactions between prenatal immune activation and Δ9-tetrahydrocannabinol exposure during adolescence in behaviours relevant to symptom dimensions of schizophrenia in rats. Prog. Neuropsychopharmacol. Biol. Psychiatry 129, 110889. 10.1016/j.pnpbp.2023.110889

Moreno-Fernández, M., Luján, V., Baliyan, S., Poza, C., Capellán, R., de Las Heras-Martínez, N., Morcillo, M.Á., Oteo, M., Ambrosio, E., Ucha, M., Higuera-Matas, A., 2025. A hidden mark of a troubled past: neuroimaging and transcriptomic analyses reveal interactive effects of maternal immune activation and adolescent THC exposure suggestive of increased neuropsychiatric risk. Biol. Psychiatry Glob. Open Sci. 5, 100452. 10.1016/j.bpsgos.2025.100452

Moser, M.B., Moser, E.I., 1998. Functional differentiation in the hippocampus. Hippocampus 8, 608–619. 10.1002/(SICI)1098-1063(1998)8:6<608::AID-HIPO3>3.0.CO;2-7

Nakagawa, K., Yoshino, H., Ogawa, Y., Yamamuro, K., Kimoto, S., Noriyama, Y., Makinodan, M., Yamashita, M., Saito, Y., Kishimoto, T., 2020. Maternal immune activation affects hippocampal excitatory and inhibitory synaptic transmission in offspring from an early developmental period to adulthood. Front. Cell. Neurosci. 14, 241. 10.3389/fncel.2020.00241

Nasyrova, R.F., Ivashchenko, D.V., Ivanov, M.V., Neznanov, N.G., 2015. Role of nitric oxide and related molecules in schizophrenia pathogenesis: biochemical, genetic and clinical aspects. Front. Physiol. 6, 139. 10.3389/fphys.2015.00139

Navarrete, F., García-Gutiérrez, M.S., Jurado-Barba, R., Rubio, G., Gasparyan, A., Austrich-Olivares, A., Manzanares, J., 2020. Endocannabinoid system components as potential biomarkers in psychiatry. Front. Psychiatry 11, 315. 10.3389/fpsyt.2020.00315

Paoletti, P., Bellone, C., Zhou, Q., 2013. NMDA receptor subunit diversity: impact on receptor properties, synaptic plasticity and disease. Nat. Rev. Neurosci. 14, 383–400. 10.1038/nrn3504

Pfaffl, M.W., 2001. A new mathematical model for relative quantification in real-time RT-PCR. Nucleic Acids Res. 29, e45. 10.1093/nar/29.9.e45

Pujante-Gil, S., Manzanedo, C., Arenas, M.C., 2021. Efecto del estrés sobre la inhibición por prepulso: revisión sistemática. Rev. Neurol. 72, 121–132. 10.33588/rn.7204.2020441

Rahman, T., Zavitsanou, K., Purves-Tyson, T., Harms, L.R., Meehan, C., Schall, U., Todd, J., Hodgson, D.M., Michie, P.T., Shannon Weickert, C., 2017. Effects of immune activation during early or late gestation on N-methyl-D-aspartate receptor measures in adult rat offspring. Front. Psychiatry 8, 77. 10.3389/fpsyt.2017.00077

Rodenas-Ruano, A., Chávez, A.E., Cossio, M.J., Castillo, P.E., Zukin, R.S., 2012. REST-dependent epigenetic remodeling promotes the developmental switch in synaptic NMDA receptors. Nat. Neurosci. 15, 1382–1390. 10.1038/nn.3214

Roura-Martínez, D., Ucha, M., Orihuel, J., Ballesteros-Yáñez, I., Castillo, C.A., Marcos, A., Ambrosio, E., Higuera-Matas, A., 2020. Central nucleus of the amygdala as a common substrate of the incubation of drug and natural reinforcer seeking. Addict. Biol. 25, e12706. 10.1111/adb.12706

Rovný, R., Marko, M., Katina, S., Murínová, J., Roháriková, V., Cimrová, B., Repiská, G., Minárik, G., Riečanský, I., 2018. Association between genetic variability of neuronal nitric oxide synthase and sensorimotor gating in humans. Nitric Oxide 80:32–36. 10.1016/j.niox.2018.08.002

Rubenstein, J.L.R., Merzenich, M.M., 2003. Model of autism: increased ratio of excitation/inhibition in key neural systems. Genes Brain Behav. 2, 255–267. 10.1034/j.1601-183x.2003.00037.x

Ruijter, J.M., Ramakers, C., Hoogaars, W.M.H., Karlen, Y., Bakker, O., van den Hoff, M.J.B., Moorman, A.F.M., 2009. Amplification efficiency: linking baseline and bias in the analysis of quantitative PCR data. Nucleic Acids Res. 37, e45. 10.1093/nar/gkp045

Santoni, M., Pistis, M., 2025. Maternal immune activation and the endocannabinoid system: focus on two-hit models of schizophrenia. Biol. Psychiatry 97, 105–115. 10.1016/j.biopsych.2024.11.015

Santos-Toscano, R., Borcel, É., Ucha, M., Orihuel, J., Capellán, R., Roura-Martínez, D., Ambrosio, E., Higuera-Matas, A., 2016. Unaltered cocaine self-administration in the prenatal LPS rat model of schizophrenia. Prog. Neuropsychopharmacol. Biol. Psychiatry 69, 38–48. 10.1016/j.pnpbp.2016.04.008

Schweizer, C., Balsiger, S., Bluethmann, H., Mansuy, I.M., Fritschy, J.M., Mohler, H., Lüscher, B., 2003. The gamma 2 subunit of GABA(A) receptors is required for maintenance of receptors at mature synapses. Mol. Cell. Neurosci. 24, 442–450. 10.1016/s1044-7431(03)00202-1

Sears, S.M.S., Hewett, S.J., 2021. Influence of glutamate and GABA transport on brain excitatory/inhibitory balance. Exp. Biol. Med. 246, 1069–1083. 10.1177/1535370221989263

Shao, Y., Cai, Y., Chen, T., Hao, K., Luo, B., Wang, X., Guo, W., Su, X., Lv, L., Yang, Y., Li, W., 2023. Impaired erythropoietin-producing hepatocellular B receptors signaling in the prefrontal cortex and hippocampus following maternal immune activation in male rats. Genes Brain Behav. 22, e12863. 10.1111/gbb.12863

Sohal, V.S., Rubenstein, J.L.R., 2019. Excitation-inhibition balance as a framework for investigating mechanisms in neuropsychiatric disorders. Mol. Psychiatry 24, 1248–1257. 10.1038/s41380-019-0426-0

Tellez-Merlo, G., Morales-Medina, J.C., Camacho-Ábrego, I., Juárez-Díaz, I., Aguilar-Alonso, P., de la Cruz, F., Iannitti, T., Flores, G., 2019. Prenatal immune challenge induces behavioral deficits, neuronal remodeling, and increases brain nitric oxide and zinc levels in the male rat offspring. Neuroscience 406, 594–605. 10.1016/j.neuroscience.2019.02.018

Tricoire, L., Vitalis, T., 2012. Neuronal nitric oxide synthase expressing neurons: a journey from birth to neuronal circuits. Front. Neural Circuits 6, 82. 10.3389/fncir.2012.00082

Uliana, D.L., Lisboa, J.R.F., Gomes, F.V., Grace, A.A., 2024. The excitatory-inhibitory balance as a target for the development of novel drugs to treat schizophrenia. Biochem. Pharmacol. 228, 116298. 10.1016/j.bcp.2024.116298

Wu, J.Y., Prentice, H., 2010. Role of taurine in the central nervous system. J. Biomed. Sci. 17 (Suppl. 1), S1. 10.1186/1423-0127-17-S1-S1

Yashiro, K., Philpot, B.D., 2008. Regulation of NMDA receptor subunit expression and its implications for LTD, LTP, and metaplasticity. Neuropharmacology 55, 1081–1094. 10.1016/j.neuropharm.2008.07.046

Zhang, J., Jing, Y., Zhang, H., Bilkey, D.K., Liu, P., 2017. Maternal immune activation leads to increased nNOS immunoreactivity in the brain of postnatal day 2 rat offspring. Synapse 72, e22011. 10.1002/syn.22011

Zhang, Z., Wang, W., Zhong, P., Liu, S.J., Long, J.Z., Zhao, L., Gao, H.Q., Cravatt, B.F., Liu, Q.S., 2015. Blockade of 2-arachidonoylglycerol hydrolysis produces antidepressant-like effects and enhances adult hippocampal neurogenesis and synaptic plasticity. Hippocampus 25, 16–26. 10.1002/hipo.22344

Zhu, Y., Wang, R., Fan, Z., Luo, D., Cai, G., Li, X., Han, J., Zhuo, L., Zhang, L., Zhang, H., Li, Y., Wu, S., 2022. Taurine alleviates chronic social defeat stress-induced depression by protecting cortical neurons from dendritic spine loss. Cell. Mol. Neurobiol. 43, 827–840. 10.1007/s10571-022-01218-3

