## Supplementary Information for "Maternal immune activation and peripubertal stress differentially disrupt glutamatergic, endocannabinoid, and neuromodulatory signalling in the adult rat dorsal hippocampus: implications for excitatory-inhibitory balance"

**Supplementary Materials and Methods**

**Table S1. List of qPCR primers used**

| Gene identification: protein (*gene*) | | Entire name | forward primer sequence | reverse primer sequence |
| --- | --- | --- | --- | --- |
| *Housekeeping gene:* | |  | |  |
|  | GAPDH *(Gapdh)* | Glyceraldehyde 3-phosphate dehydrogenase | 5'-tccctgttctagagacag-3' | 5'-ccactttgtcacaagaga-3' |
| *Genes of glutamatergic ionotropic receptors subunits:* | |  | |  |
|  | NR1 *(Grin1)* | NMDA receptor subunit 1 | 5'-aacctgcagaaccgcaag-3' | 5'-gcttgatgagcaggtctatgc-3' |
|  | NR2A *(Grin2a)* | NMDA receptor subunit 2A | 5'-tgtgaagaaatgctgcaagg-3' | 5'-gaacgctcctcattgatggt-3' |
|  | NR2B *(Grin2b)* | NMDA receptor subunit 2B | 5'-caatggcagcacagagagga-3' | 5'-agcaatgccatagccggtag-3' |
|  | GluA1 *(Gria1)* | AMPA receptor subunit 1 | 5'-agaggctggtggtggttgact-3' | 5'-accctggtatggtctcggga-3' |
|  | GluA2 *(Gria2)* | AMPA receptor subunit 2 | 5'-ggcgtgtaatcctggactgt-3' | 5'-acaccagggaatcgtcgtag-3' |
| *Genes of GABAergic ionotropic receptors subunits:* | |  | |  |
|  | GABA_A_γ2 *(Gabrg2)* | GABA_A_ receptor subunit gamma 2 | 5'-cggaaaccaagcaaggataa-3' | 5'-acagtccttgccatccaaac-3' |
| *Genes of the endocannabinoid system:* | |  | |  |
|  | CB1 *(Cnr1)* | cannabinoid receptor 1 | 5'-gtcgatcctagatggccttgc-3' | 5'-gtcattcgagcccacgtagag-3' |
|  | NAPE-PLD *(Napepld)* | *N*-acylphosphatidylethanolamine phospholipase D | 5'-accaacatgctgacccagag-3' | 5'-atcgtgactctccgtgcttc-3' |
|  | FAAH *(Faah)* | fatty acid amidohydrolase | 5'-gttacagagtggagagctgtcc-3' | 5'-gtctcacagtcggtcagatagg-3' |
|  | DAGLα *(Dagla)* | diacylglycerol lipase alpha | 5'-ctttgctgaatttttccgtgacc-3' | 5'-ttgtttgcctcatccagcac-3' |
|  | MAGL *(Mgll)* | monoacylglycerol lipase | 5'-ctacctgctcatggaatc-3' | 5'-gacacccacgtatttatttc-3' |
| *Genes of the nitric oxide system:* | |  | |  |
| nNOS (*Nos1*) | | neuronal nitric oxide synthase 5'-tgagtccatcgccttcatcg-3' | | 5'-acaccagggaatcgtcgtag-3' |

**Supplementary results**

| **PPI condition** | **Prenatal Treatment** | **Postnatal Treatment** | **Mean** | **Standard Deviation** | **N** |
| --- | --- | --- | --- | --- | --- |
| PPI12_120 | Saline | No stress | 32.179 | 14.497 | 9 |
|  |  | Stress | 46.174 | 9.656 | 9 |
|  |  | Total | 39.176 | 13.951 | 18 |
|  | LPS | No stress | 37.450 | 18.541 | 6 |
|  |  | Stress | 35.436 | 8.011 | 7 |
|  |  | Total | 36.365 | 13.282 | 13 |
|  | Total | No stress | 34.287 | 15.812 | 15 |
|  |  | Stress | 41.476 | 10.279 | 16 |
|  |  | Total | 37.997 | 13.521 | 31 |
| PPI4_120 | Saline | No stress | 36.281 | 20.713 | 7 |
|  |  | Stress | 29.319 | 12.829 | 8 |
|  |  | Total | 32.568 | 16.706 | 15 |
|  | LPS | No stress | 30.381 | 11.356 | 6 |
|  |  | Stress | 26.065 | 9.966 | 6 |
|  |  | Total | 28.223 | 10.432 | 12 |
|  | Total | No stress | 33.558 | 16.662 | 13 |
|  |  | Stress | 27.925 | 11.384 | 14 |
|  |  | Total | 30.637 | 14.183 | 27 |
| PPI12_30 | Saline | No stress | 31.379 | 17.054 | 9 |
|  |  | Stress | 46.931 | 18.130 | 10 |
|  |  | Total | 39.564 | 18.901 | 19 |
|  | LPS | No stress | 34.976 | 22.768 | 8 |
|  |  | Stress | 38.086 | 16.116 | 7 |
|  |  | Total | 36.427 | 19.315 | 15 |
|  | Total | No stress | 33.072 | 19.381 | 17 |
|  |  | Stress | 43.289 | 17.390 | 17 |
|  |  | Total | 38.180 | 18.858 | 34 |
| PPI4_30 | Saline | No stress | 13.415 | 9.013 | 4 |
|  |  | Stress | 29.201 | 15.356 | 7 |
|  |  | Total | 23.461 | 15.142 | 11 |
|  | LPS | No stress | 8.845 | 9.673 | 5 |
|  |  | Stress | 20.525 | 11.255 | 5 |
|  |  | Total | 14.685 | 11.652 | 10 |
|  | Total | No stress | 10.876 | 9.113 | 9 |
|  |  | Stress | 25.586 | 13.952 | 12 |
|  |  | Total | 19.282 | 13.997 | 21 |

**Supplementary Table S2. Descriptive statistics of prepulse inhibition (PPI) per group and PPI condition. Negative PPI values (prepulse facilitation) were excluded prior to analysis. SAL: saline; LPS: lipopolysaccharide; NS: no stress; S: stress.**

| **Effect** | **PPI condition** | **F-value and degrees of freedom** | **p-value** |
| --- | --- | --- | --- |
| MIA | PPI12_120 | F (1,27) = 0,335 | 0.568 |
|  | PPI4_120 | F (1,23) = 0,659 | 0.425 |
|  | PPI12_30 | F (1,30) = 0,165 | 0.688 |
|  | PPI4_30 | F (1,17) = 1,481 | 0.240 |
| PUS | PPI12_120 | F (1,23) = 1,000 | 0.328 |
|  | PPI4_120 | F (1,28) = 0,012 | 0.913 |
|  | PPI12_30 | F (1,30) = 2,083 | 0.159 |
|  | **PPI4_30** | **F (1,17) = 6,369** | **0.022*** |
| MIA x PUS interaction | PPI12_120 | F (1,28) = 2,872 | 0.102 |
|  | PPI4_120 | F (1,23) = 0,055 | 0.817 |
|  | PPI12_30 | F (1,30) = 0,926 | 0.344 |
|  | PPI4_30 | F (1,17) = 0,142 | 0.711 |

**Supplementary Table S3. Two-way ANOVA (MIA × PUS) results for each PPI condition. Bold entries indicate significant effects (p < 0.05). * p < 0.05.**

| **Effect** | **beta** | **SE** | **z-value** | **p-value** | **Sig.** |
| --- | --- | --- | --- | --- | --- |
| **Intercept (SAL/NS/30ms)** | **31.38** | 5.11 | 6.14 | **<0.001** | ******* |
| MIA (LPS vs SAL) | 9.77 | 7.72 | 1.27 | 0.206 | ns |
| **PUS (Stress vs No stress)** | **15.55** | 7.04 | 2.21 | **0.027** | ***** |
| ISI (120ms vs 30ms) | 0.80 | 5.49 | 0.15 | 0.884 | ns |
| MIA x PUS | -18.61 | 10.80 | -1.72 | 0.085 | (+) |
| MIA x ISI | -5.82 | 8.59 | -0.68 | 0.498 | ns |
| PUS x ISI | -3.36 | 7.74 | -0.43 | 0.664 | ns |
| MIA x PUS x ISI | 5.73 | 11.96 | 0.48 | 0.632 | ns |

**Supplementary Table S4. Linear mixed model results — 12 dB prepulse trials (PPI ~ MIA × PUS × ISI + (1|subject)). N = 64 observations, 31 subjects. REML estimation.** (+) p < 0.10 (trend); * p < 0.05; ** p < 0.01; *** p < 0.001; ns: not significant. beta = fixed effect estimate; SE = standard error. Reference category in all models: SAL (saline), NS (no stress), with 30ms ISI or 4dB as applicable. MIA: maternal immune activation (LPS vs saline, GD15-16); PUS: peripubertal unpredictable stress (PND28-38); ISI: stimulus onset asynchrony; dB: prepulse intensity above background noise.

| **Effect** | **beta** | **SE** | **z-value** | **p-value** | **Sig.** |
| --- | --- | --- | --- | --- | --- |
| **Intercept (SAL/NS/30ms)** | **15.58** | 7.17 | 2.17 | **0.030** | ***** |
| MIA (LPS vs SAL) | -7.25 | 9.55 | -0.76 | 0.448 | ns |
| PUS (Stress vs No stress) | 13.76 | 8.77 | 1.57 | 0.117 | ns |
| **ISI (120ms vs 30ms)** | **20.70** | 8.17 | 2.53 | **0.011** | ***** |
| MIA x PUS | -1.89 | 12.30 | -0.15 | 0.878 | ns |
| MIA x ISI | 1.35 | 11.17 | 0.12 | 0.904 | ns |
| **PUS x ISI** | **-20.71** | 10.29 | -2.01 | **0.044** | ***** |
| MIA x PUS x ISI | 4.50 | 14.79 | 0.30 | 0.761 | ns |

**Supplementary Table S5. Linear mixed model results — 4 dB prepulse trials (PPI ~ MIA × PUS × ISI + (1|subject)). N = 48 observations, 28 subjects. REML estimation.** (+) p < 0.10 (trend); * p < 0.05; ** p < 0.01; *** p < 0.001; ns: not significant. beta = fixed effect estimate; SE = standard error. Reference category in all models: SAL (saline), NS (no stress), with 30ms ISI or 4dB as applicable. MIA: maternal immune activation (LPS vs saline, GD15-16); PUS: peripubertal unpredictable stress (PND28-38); ISI: stimulus onset asynchrony; dB: prepulse intensity above background noise.

| **Effect** | **beta** | **SE** | **z-value** | **p-value** | **Sig.** |
| --- | --- | --- | --- | --- | --- |
| Intercept (SAL/NS/4dB) | 12.62 | 7.42 | 1.70 | 0.089 | (+) |
| MIA (LPS vs SAL) | -5.69 | 10.07 | -0.57 | 0.572 | ns |
| PUS (Stress vs No stress) | 14.98 | 9.38 | 1.60 | 0.110 | ns |
| **dB (12dB vs 4dB)** | **18.76** | 7.78 | 2.41 | **0.016** | ***** |
| MIA x PUS | -2.09 | 13.45 | -0.16 | 0.876 | ns |
| MIA x dB | 15.46 | 10.66 | 1.45 | 0.147 | ns |
| PUS x dB | 0.57 | 9.91 | 0.06 | 0.954 | ns |
| MIA x PUS x dB | -16.52 | 14.33 | -1.15 | 0.249 | ns |

**Supplementary Table S6. Linear mixed model results — ISI 30 ms trials (PPI ~ MIA × PUS × dB + (1|subject)). N = 54 observations, 30 subjects. REML estimation.** (+) p < 0.10 (trend); * p < 0.05; ** p < 0.01; *** p < 0.001; ns: not significant. beta = fixed effect estimate; SE = standard error. Reference category in all models: SAL (saline), NS (no stress), with 30ms ISI or 4dB as applicable. MIA: maternal immune activation (LPS vs saline, GD15-16); PUS: peripubertal unpredictable stress (PND28-38); ISI: stimulus onset asynchrony; dB: prepulse intensity above background noise.

| **Effect** | **beta** | **SE** | **z-value** | **p-value** | **Sig.** |
| --- | --- | --- | --- | --- | --- |
| **Intercept (SAL/NS/4dB)** | **36.16** | 4.96 | 7.30 | **<0.001** | ******* |
| MIA (LPS vs SAL) | -5.78 | 7.45 | -0.78 | 0.438 | ns |
| PUS (Stress vs No stress) | -7.63 | 6.85 | -1.11 | 0.266 | ns |
| dB (12dB vs 4dB) | -3.98 | 4.67 | -0.85 | 0.394 | ns |
| MIA x PUS | 2.40 | 10.35 | 0.23 | 0.817 | ns |
| MIA x dB | 11.05 | 6.97 | 1.59 | 0.113 | ns |
| **PUS x dB** | **21.62** | 6.44 | 3.36 | **<0.001** | ******* |
| MIA x PUS x dB | -18.41 | 9.70 | -1.90 | 0.058 | (+) |

**Supplementary Table S7. Linear mixed model results — ISI 120 ms trials (PPI ~ MIA × PUS × dB + (1|subject)). N = 58 observations, 31 subjects. REML estimation.** (+) p < 0.10 (trend); * p < 0.05; ** p < 0.01; *** p < 0.001; ns: not significant. beta = fixed effect estimate; SE = standard error. Reference category in all models: SAL (saline), NS (no stress), with 30ms ISI or 4dB as applicable. MIA: maternal immune activation (LPS vs saline, GD15-16); PUS: peripubertal unpredictable stress (PND28-38); ISI: stimulus onset asynchrony; dB: prepulse intensity above background noise.

| **Prenatal treatment** | **Postnatal treatment** | **Mean** | **Standard**  **Deviation** | **N** |
| --- | --- | --- | --- | --- |
| **GABA** | | | | |
| Saline | No stress | 53.358 | 10.401 | 9 |
|  | Stress | 46.025 | 8.250 | 10 |
|  | Total | 49.499 | 9.813 | 19 |
| LPS | No stress | 50.875 | 5.209 | 8 |
|  | Stress | 52.644 | 7.512 | 7 |
|  | Total | 51.700 | 6.212 | 15 |
| Total | No stress | 52.190 | 8.224 | 17 |
|  | Stress | 48.751 | 8.409 | 17 |
|  | Total | 50.470 | 8.374 | 34 |
| **L-Glutamine** | | | | |
| Saline | No stress | 177.123 | 43.586 | 9 |
|  | Stress | 177.437 | 45.929 | 10 |
|  | Total | 177.288 | 43.578 | 19 |
| LPS | No stress | 181.492 | 20.189 | 8 |
|  | Stress | 190.169 | 22.525 | 7 |
|  | Total | 185.541 | 21.007 | 15 |
| Total | No stress | 179.179 | 33.664 | 17 |
|  | Stress | 182.929 | 37.663 | 17 |
|  | Total | 180.929 | 35.219 | 34 |
| **Glycine** | | | | |
| Saline | No stress | 54.090 | 10.924 | 9 |
|  | Stress | 53.220 | 4.643 | 8 |
|  | Total | 53.680 | 8.325 | 17 |
| LPS | No stress | 59.728 | 13.174 | 8 |
|  | Stress | 62.917 | 6.417 | 7 |
|  | Total | 61.216 | 10.351 | 15 |
| Total | No stress | 56.743 | 12.001 | 17 |
|  | Stress | 57.745 | 7.315 | 15 |
|  | Total | 57.213 | 9.937 | 32 |
| **L-Serine** | | | | |
| Saline | No stress | 206.531 | 64.518 | 9 |
|  | Stress | 197.425 | 36.506 | 10 |
|  | Total | 201.738 | 50.380 | 19 |
| LPS | No stress | 181.331 | 42.630 | 8 |
|  | Stress | 230.031 | 34.687 | 7 |
|  | Total | 204.057 | 45.351 | 15 |
| Total | No stress | 194.672 | 55.176 | 17 |
|  | Stress | 210.851 | 38.398 | 17 |
|  | Total | 202.761 | 47.522 | 34 |
| **Taurine** | | | | |
| Saline | No stress | 99.755 | 21.550 | 9 |
|  | Stress | 90.740 | 17.238 | 10 |
|  | Total | 95.010 | 19.400 | 19 |
| LPS | No stress | 90.697 | 8.674 | 8 |
|  | Stress | 107.300 | 9.530 | 7 |
|  | Total | 98.445 | 12.250 | 15 |
| Total | No stress | 95.492 | 16.485 | 17 |
|  | Stress | 97.559 | 16.485 | 17 |
|  | Total | 96.526 | 16.491 | 34 |
| **L-Glutamate** | | | | |
| Saline | No stress | 446.741 | 68.837 | 8 |
|  | Stress | 436.707 | 72.206 | 10 |
|  | Total | 441.166 | 68.831 | 18 |
| LPS | No stress | 430.653 | 48.537 | 8 |
|  | Stress | 459.634 | 60.194 | 7 |
|  | Total | 444.177 | 54.357 | 15 |
| Total | No stress | 438.697 | 58.135 | 16 |
|  | Stress | 446.147 | 66.533 | 17 |
|  | Total | 442. 535 | 61.741 | 33 |
| **Aspartate** | | | | |
| Saline | No stress | 190.587 | 33.420 | 8 |
|  | Stress | 186.993 | 40.858 | 10 |
|  | Total | 188.590 | 36.702 | 18 |
| LPS | No stress | 168.040 | 18.577 | 8 |
|  | Stress | 204.981 | 16.704 | 7 |
|  | Total | 185.279 | 25.613 | 15 |
| Total | No stress | 179.313 | 28.598 | 16 |
|  | Stress | 194.400 | 33.570 | 17 |
|  | Total | 187.085 | 31.708 | 33 |
| **L-Gln/L-Glu ratio** | | | | |
| Saline | No stress | 0.382 | 0.069 | 9 |
|  | Stress | 0.403 | 0.048 | 10 |
|  | Total | 0.393 | 0.058 | 19 |
| LPS | No stress | 0.425 | 0.062 | 8 |
|  | Stress | 0.417 | 0.045 | 7 |
|  | Total | 0.421 | 0.053 | 15 |
| Total | No stress | 0.402 | 0.067 | 17 |
|  | Stress | 0.409 | 0.046 | 17 |
|  | Total | 0.405 | 0.057 | 34 |

**Supplementary Table S8. Descriptive statistics for hippocampal amino acid concentrations (µM) measured by CE-LIF.**

| **Effect and Treatment** | **F-value and degrees of freedom** | **p-value** |
| --- | --- | --- |
| GABA | | |
| MIA | F (1,30) = 0.533 | 0.471 |
| PUS | F (1,30) = 0.996 | 0.334 |
| MIA x PUS interaction | F (1,30) = 2.585 | 0.118 |
| L-Glutamine | | |
| MIA | F (1,30) = 0.457 | 0.504 |
| PUS | F (1,30) = 0.126 | 0.725 |
| MIA x PUS interaction | F (1,30) = 0.109 | 0.743 |
| Glycine | | |
| MIA | **F (1,28) = 5.089** | **0.032*** |
| PUS | F (1,28) = 0.116 | 0.736 |
| MIA x PUS interaction | F (1,28) = 0.357 | 0.555 |
| L-Serine | | |
| MIA | F (1,30) = 0.053 | 0.820 |
| PUS | F (1,30) = 1.505 | 0.229 |
| MIA x PUS interaction | F (1,30) = 3.208 | 0.083 |
| Taurine | | |
| MIA | F (1,30) = 0.472 | 0.497 |
| PUS | F (1,30) = 0.483 | 0.492 |
| MIA x PUS interaction | **F (1,30) = 5.510** | **0.026*** |
| L-Glutamate | | |
| MIA | F (1,29) = 0.023 | 0.880 |
| PUS | F (1,29) = 0.179 | 0.676 |
| MIA x PUS interaction | F (1,29) = 0.757 | 0.391 |
| Aspartate | | |
| MIA | F (1,29) = 0.045 | 0.833 |
| PUS | F (1,29) = 2.429 | 0.130 |
| MIA x PUS interaction | F (1,29) = 3.590 | 0.068 |
| L-Glutamine/L-Glutamate ratio | | |
| MIA | F (1,30) = 2.072 | 0.160 |
| PUS | F (1,30) = 0.100 | 0.754 |
| MIA x PUS interaction | F (1,30) = 0.567 | 0.457 |

**Table S9. ANOVA table for each metabolite analysed by capillary electrophoresis for each condition and their interactions.**

| **Prenatal treatment** | **Postnatal treatment** | **Mean** | **Standard Deviation** | **N** |
| --- | --- | --- | --- | --- |
| *Gria1* | | | | |
| Saline | No stress | 1.000 | 0.375 | 10 |
|  | Stress | 1.165 | 0.624 | 10 |
|  | Total | 1.082 | 0.508 | 20 |
| LPS | No stress | 1.158 | 0.475 | 7 |
|  | Stress | 1.122 | 0.276 | 8 |
|  | Total | 1.145 | 0.367 | 15 |
| Total | No stress | 1.065 | 0.412 | 17 |
|  | Stress | 1.151 | 0.487 | 18 |
|  | total | 1.109 | 0.448 | 35 |
| *Gria2* | | | | |
| Saline | No stress | 1.000 | 0.528 | 10 |
|  | Stress | 1.095 | 0.499 | 9 |
|  | Total | 1.116 | 0.503 | 19 |
| LPS | No stress | 1.090 | 0.411 | 7 |
|  | Stress | 1.148 | 0.206 | 6 |
|  | Total | 1.169 | 0.321 | 13 |
| Total | No stress | 1.037 | 0.472 | 17 |
|  | Stress | 1.116 | 0.398 | 15 |
|  | Total | 1.074 | 0.433 | 32 |
| *Grm1* | | | | |
| Saline | No stress | 1.000 | 0.420 | 10 |
|  | Stress | 0.980 | 0.287 | 10 |
|  | Total | 0.990 | 0.350 | 20 |
| LPS | No stress | 1.026 | 0.467 | 7 |
|  | Stress | 1.213 | 0.287 | 8 |
|  | Total | 1.126 | 0.380 | 15 |
| Total | No stress | 1.011 | 0.426 | 17 |
|  | Stress | 1.084 | 0.303 | 18 |
|  | Total | 1.048 | 0.364 | 35 |
| *Grin1* | | | | |
| Saline | No stress | 1.000 | 0.623 | 9 |
|  | Stress | 1.132 | 0.685 | 8 |
|  | Total | 1.064 | 0.636 | 17 |
| LPS | No stress | 0.573 | 0.430 | 8 |
|  | Stress | 0.926 | 0.300 | 7 |
|  | Total | 0.738 | 0.405 | 15 |
| Total | No stress | 0.802 | 0.567 | 17 |
|  | Stress | 1.036 | 0.534 | 15 |
|  | Total | 0.911 | 0.557 | 32 |
| *Grin2A* | | | | |
| Saline | No stress | 1.000 | 0.378 | 10 |
|  | Stress | 1.002 | 0.334 | 9 |
|  | Total | 1.001 | 0.348 | 19 |
| LPS | No stress | 1.105 | 0.487 | 7 |
|  | Stress | 1.273 | 0.472 | 8 |
|  | Total | 1.194 | 0.470 | 15 |
| Total | No stress | 1.043 | 0.415 | 17 |
|  | Stress | 1.130 | 0.415 | 17 |
|  | Total | 1.086 | 0.411 | 34 |
| *Grin2B* | | | | |
| Saline | No stress | 1.000 | 0.429 | 10 |
|  | Stress | 1.337 | 0.6161 | 10 |
|  | Total | 1.169 | 0.545 | 20 |
| LPS | No stress | 1.069 | 0.482 | 7 |
|  | Stress | 1.373 | 0.436 | 8 |
|  | Total | 1.231 | 0.468 | 15 |
| Total | No stress | 1.029 | 0.438 | 17 |
|  | Stress | 1.353 | 0.529 | 18 |
|  | Total | 1.195 | 0.507 | 35 |
| *Gabrg2* | | | | |
| Saline | No stress | 1,000 | 0,311 | 10 |
|  | Stress | 0,919 | 0,265 | 10 |
|  | Total | 0,959 | 0,284 | 20 |
| LPS | No stress | 0,870 | 0,247 | 7 |
|  | Stress | 0,815 | 0,141 | 7 |
|  | Total | 0,842 | 0,195 | 14 |
| Total | No stress | 0,947 | 0,285 | 17 |
|  | Stress | 0,876 | 0,223 | 17 |
|  | Total | 0,911 | 0,255 | 34 |
| *Gria2/Gria1 ratio* | | | | |
| Saline | No stress | 1,475 | 2,059 | 10 |
|  | Stress | 1,215 | 0,784 | 10 |
|  | Total | 1,345 | 1,523 | 20 |
| LPS | No stress | 1,786 | 2,661 | 7 |
|  | Stress | 1,052 | 0,290 | 7 |
|  | Total | 1,419 | 1,858 | 14 |
| Total | No stress | 1,603 | 2,251 | 17 |
|  | Stress | 1,148 | 0,620 | 17 |
|  | Total | 1,375 | 1,642 | 34 |
| *Grin2A/Grin2B ratio* | | | | |
| Saline | No stress | 1,035 | 0,133 | 10 |
|  | Stress | 0,854 | 0,091 | 9 |
|  | Total | 0,949 | 0,145 | 19 |
| LPS | No stress | 1,022 | 0,085 | 7 |
|  | Stress | 0,919 | 0,132 | 8 |
|  | Total | 0,967 | 0,121 | 15 |
| Total | No stress | 1,030 | 0,113 | 17 |
|  | Stress | 0,885 | 0,114 | 17 |
|  | Total | 0,957 | 0,134 | 34 |

**Table S10. ANOVA table for gene expression data for each of the genes analysed and their ratios related to the glutamatergic and GABAergic system.**

| **Prenatal treatment** | **Postnatal treatment** | **Mean** | **Standard Deviation** | **N** |
| --- | --- | --- | --- | --- |
| *Cnr1* | | | | |
| Saline | No stress | 1,000 | 0,344 | 10 |
|  | Stress | 0,940 | 0,345 | 10 |
|  | Total | 0,970 | 0,337 | 20 |
| LPS | No stress | 1,000 | 0,400 | 7 |
|  | Stress | 1,008 | 0,216 | 8 |
|  | Total | 1,004 | 0,301 | 15 |
| Total | No stress | 1,000 | 0,354 | 17 |
|  | Stress | 0,970 | 0,289 | 18 |
|  | Total | 0,985 | 0,318 | 35 |
| *Dagla* | | | | |
| Saline | No stress | 1,000 | 0,509 | 10 |
|  | Stress | 0,968 | 0,204 | 10 |
|  | Total | 0,984 | 0,378 | 20 |
| LPS | No stress | 0,573 | 0,184 | 7 |
|  | Stress | 0,992 | 0,253 | 8 |
|  | Total | 0,797 | 0,306 | 15 |
| Total | No stress | 0,824 | 0,453 | 17 |
|  | Stress | 0,979 | 0,221 | 18 |
|  | Total | 0,904 | 0,356 | 35 |
| *Mgll* | | | | |
| Saline | No stress | 1,000 | 0,304 | 10 |
|  | Stress | 1,106 | 0,359 | 10 |
|  | Total | 1,053 | 0,280 | 20 |
| LPS | No stress | 0,813 | 0,126 | 6 |
|  | Stress | 1,113 | 0,320 | 8 |
|  | Total | 0,984 | 0,291 | 14 |
| Total | No stress | 0,930 | 0,264 | 16 |
|  | Stress | 1,109 | 0,279 | 18 |
|  | Total | 1,025 | 0,283 | 34 |
| *Faah* | | | | |
| Saline | No stress | 0,770 | 0,412 | 9 |
|  | Stress | 1,066 | 0,574 | 10 |
|  | Total | 0,926 | 0,513 | 19 |
| LPS | No stress | 0,709 | 0,565 | 7 |
|  | Stress | 0,824 | 0,398 | 8 |
|  | Total | 0,770 | 0,469 | 15 |
| Total | No stress | 0,743 | 0,468 | 16 |
|  | Stress | 0,958 | 0,505 | 18 |
|  | Total | 0,857 | 0,493 | 34 |
| *Napepdl* | | | | |
| Saline | No stress | 0,887 | 0,239 | 9 |
|  | Stress | 0,862 | 0,243 | 10 |
|  | Total | 0,874 | 0,235 | 19 |
| LPS | No stress | 0,786 | 0,168 | 7 |
|  | Stress | 0,928 | 0,380 | 8 |
|  | Total | 0,862 | 0,300 | 15 |
| Total | No stress | 0,843 | 0,211 | 16 |
|  | Stress | 0,891 | 0,303 | 18 |
|  | Total | 0,868 | 0,261 | 34 |
| *Faah/Napepdl ratio* | | | | |
| Saline | No stress | 0,866 | 0,496 | 8 |
|  | Stress | 1,421 | 1,029 | 10 |
|  | Total | 1,175 | 0,862 | 18 |
| LPS | No stress | 1,020 | 1,001 | 7 |
|  | Stress | 1,009 | 0,637 | 8 |
|  | Total | 1,014 | 0,795 | 15 |
| Total | No stress | 0,928 | 0,747 | 15 |
|  | Stress | 1,238 | 0,879 | 18 |
|  | Total | 1,102 | 0,823 | 33 |
| *Mgll/Dagl ratio* | | | | |
| Saline | No stress | 1,173 | 0,495 | 10 |
|  | Stress | 1,146 | 0,171 | 10 |
|  | Total | 1,159 | 0,361 | 20 |
| LPS | No stress | 1,290 | 0,142 | 6 |
|  | Stress | 1,130 | 0,187 | 8 |
|  | Total | 1,199 | 0,183 | 14 |
| Total | No stress | 1,217 | 0,396 | 16 |
|  | Stress | 1,139 | 0,174 | 18 |
|  | Total | 1,176 | 0,298 | 34 |

**Table S11. ANOVA table for gene expression data for each of the genes analysed and their ratios related to the endocannabinoid system.**

| **Prenatal treatment** | **Postnatal treatment** | **Mean** | **Standard Deviation** | **N** |
| --- | --- | --- | --- | --- |
| *Nos1* | | | | |
| Saline | No stress | 0.889 | 0.250 | 10 |
|  | Stress | 1.242 | 0.460 | 9 |
|  | Total | 1.061 | 0.396 | 19 |
| LPS | No stress | 0.886 | 0.295 | 6 |
|  | Stress | 0.920 | 0.354 | 7 |
|  | Total | 0.904 | 0.315 | 13 |
| Total | No stress | 0.894 | 0.258 | 16 |
|  | Stress | 1.101 | 0.436 | 16 |
|  | Total | 0.997 | 0.368 | 32 |

| **Effect and Treatment** | **F-value and degrees of freedom** | **p-value** |
| --- | --- | --- |
| *Gria1* |  |  |
| MIA | F (1,31) = 0.158 | 0.693 |
| PUS | F (1,31) = 0.195 | 0.662 |
| MIA x PUS interaction | F (1,31) = 0.356 | 0.555 |
| *Gria2* |  |  |
| MIA | F (1,30) = 0.192 | 0.664 |
| PUS | F (1,30) = 0.311 | 0.581 |
| MIA x PUS interaction | F (1,30) = 0.024 | 0.879 |
| *Grm1* |  |  |
| MIA | F (1,31) = 1.053 | 0.313 |
| PUS | F (1,31) = 0.441 | 0.512 |
| MIA x PUS interaction | F (1,31) = 0.671 | 0.419 |
| *Grin1* |  |  |
| MIA | F (1,26) = 1.315 | 0.262 |
| PUS | F (1,26) = 0.563 | 0.460 |
| MIA x PUS interaction | F (1,26) = 0.120 | 0.915 |
| *Grina2* |  |  |
| MIA | F (1,30) = 0.201 | 0.201 |
| PUS | F (1,30) = 0.354 | 0.556 |
| MIA x PUS interaction | F (1,30) = 0.334 | 0.567 |
| *Grin2b* |  |  |
| MIA | F (1,31) = 0.094 | 0.761 |

| PUS | F (1,31) = 3.486 | 0.071 |
| --- | --- | --- |
| MIA x PUS interaction | F (1,31) = 0.009 | 0.923 |
| *Gabrg2* |  |  |
| MIA | F (1,30) = 1.705 | 0.202 |
| PUS | F (1,30) = 0.585 | 0.450 |
| MIA x PUS interaction | F (1,30) = 0.021 | 0.886 |
| *Grin2a*/*Grn2b* ratio |  |  |
| MIA | F (1,30) = 0.434 | 0.515 |
| **PUS** | **F (1,30) = 12.875** | **0.001*** |
| MIA x PUS interaction | F (1,30) = 0.981 | 0.330 |
| *Gria2*/*Gria1* ratio |  |  |
| MIA | F (1,30) = 0.016 | 0.900 |
| PUS | F (1,30) = 0.704 | 0.408 |
| MIA x PUS interaction | F (1,30) = 0.160 | 0.692 |

**Supplementary Table S12. Two-way ANOVA (MIA × PUS) results for glutamatergic and GABAergic gene expression (RT-qPCR). Bold: significant. * p < 0.05.**

| **Gene** | **F-value and degrees of freedom** | **p-value** |
| --- | --- | --- |
| *Cnr1* |  |  |
| MIA | F (1,31) = 0.090 | 0.767 |
| PUS | F (1,31) = 0.052 | 0.820 |
| MIA x PUS interaction | F (1,31) = 0.092 | 0.763 |
| *Dagla* |  |  |
| MIA | F (1,31) = 3.198 | 0.083 |
| PUS | F (1,31) = 2.954 | 0.096 |
| **MIA x PUS interaction** | **F (1,31) = 4.026** | **0.054** |
| *Mgll* |  |  |
| MIA | F (1,30) = 0.890 | 0.353 |
| **PUS** | **F (1,30) = 4.494** | **0.042*** |
| MIA x PUS interaction | F (1,30) = 1.024 | 0.320 |
| *Napepld* |  |  |
| MIA | F (1,30) = 0.035 | 0.854 |
| PUS | F (1,30) = 0.396 | 0.534 |
| MIA x PUS interaction | F (1,30) = 0.816 | 0.374 |
| *Faah* |  |  |
| MIA | F (1,30) = 0.781 | 0.384 |
| PUS | F (1,30) = 1.440 | 0.240 |
| MIA x PUS interaction | F (1,30) = 0.282 | 0.559 |
| *Mgll*/*Dagla* ratio |  |  |
| MIA | F (1,30) = 0.221 | 0.642 |
| PUS | F (1,30) = 0.755 | 0.392 |
| MIA x PUS interaction | F (1,30) = 0.380 | 0.542 |
| *Napepld*/*Faah* ratio |  |  |
| MIA | F (1,29) = 0.194 | 0.663 |
| PUS | F (1,29) = 0.867 | 0.359 |
| MIA x PUS interaction | F (1,29) = 0.939 | 0.341 |

**Supplementary Table S13. Two-way ANOVA (MIA × PUS) results for endocannabinoid system gene expression (RT-qPCR). Bold: significant. * p < 0.05.**

| **Gene** | **F-value and degrees of freedom** | **p-value** |
| --- | --- | --- |
| *Nos1* |  |  |
| MIA | F (1,28) = 1.746 | 0.197 |
| PUS | F (1,28) = 2.223 | 0.147 |
| MIA x PUS interaction | F (1,28) = 1.492 | 0.232 |

**Supplementary Table S14. Two-way ANOVA (MIA × PUS) results for nitric oxide synthase gene expression (RT-qPCR).**

| **Protein** | **F-value and degrees of freedom** | **p-value** |
| --- | --- | --- |
| GLUN1 |  |  |
| **MIA** | **F (1,29) = 5.155** | **0.031*** |
| PUS | F (1,29) = 0.474 | 0.497 |
| MIA x PUS interaction | F (1,29) = 1.183 | 0.286 |
| NOS1 |  |  |
| MIA | F (1,29) = 0.352 | 0.557 |
| PUS | F (1,29) = 0.470 | 0.499 |
| MIA x PUS interaction | F (1,29) = 0.148 | 0.704 |

**Supplementary Table S15. Two-way ANOVA (MIA × PUS) results for GluN1 and nNOS protein expression (Western blot). Bold: significant. * p < 0.05.**

**Figure S1**

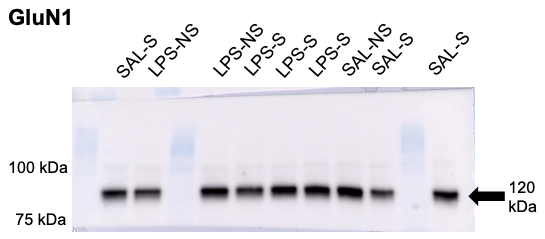

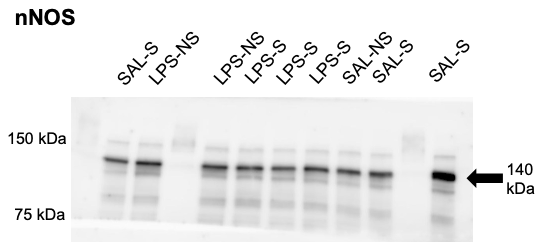

**Supplementary Figure S1 Representative images of western blots for GluN1 and nNOS proteins.**
